# Sensory neuron population expansion enhances odor tracking without sensitizing projection neurons

**DOI:** 10.1101/2023.09.15.556782

**Authors:** Suguru Takagi, Gizem Sancer, Liliane Abuin, S. David Stupski, J. Roman Arguello, Lucia L. Prieto-Godino, David L. Stern, Steeve Cruchet, Raquel Álvarez-Ocaña, Carl F. R. Wienecke, Floris van Breugel, James M. Jeanne, Thomas O. Auer, Richard Benton

**Author notes:** Department of Neurobiology, Harvard Medical School, Cambridge, Massachusetts, USA. Department of Biology, University of Fribourg, Fribourg, Switzerland (Thomas O. Auer). These authors contributed equally to this work (Suguru Takagi, Thomas O. Auer). These authors contributed equally to this work (Gizem Sancer, Liliane Abuin, S. David Stupski). Joint senior authors (Thomas O. Auer, Richard Benton).

## Abstract

The evolutionary expansion of sensory neuron populations detecting important environmental cues is widespread, but functionally enigmatic. We investigated this phenomenon through comparison of homologous neural pathways of *Drosophila melanogaster* and its close relative *Drosophila sechellia*, an extreme specialist for *Morinda citrifolia* noni fruit. *D. sechellia* has evolved species-specific expansions in select, noni-detecting olfactory sensory neuron (OSN) populations, through multigenic changes. Activation and inhibition of defined proportions of neurons demonstrate that OSN population increases contribute to stronger, more persistent, noni-odor tracking behavior. These sensory neuron expansions result in increased synaptic connections with their projection neuron (PN) partners, which are conserved in number between species. Surprisingly, having more OSNs does not lead to greater odor-evoked PN sensitivity or reliability. Rather, pathways with increased sensory pooling exhibit reduced PN adaptation, likely through weakened lateral inhibition. Our work reveals an unexpected functional impact of sensory neuron expansions to explain ecologically-relevant, species-specific behavior.

## Introduction

Brains display incredible diversity in neuron number between animal species (Godfrey and Gronenberg, 2019; Herculano-Houzel, 2011; Williams and Herrup, 1988). Increases in the number of neurons during evolution occur not only through the emergence of new cell types but also through expansion of pre-existing neuronal populations (Arendt et al., 2019; Roberts et al., 2022). Of the latter phenomenon, some of the most spectacular examples are found in sensory systems: the snout of the star-nosed mole (*Condylura cristata*) has ∼25,000 mechanosensory organs, several fold more than other mole species (Catania, 1995). Male (but not female) moths can have tens of thousands of neurons detecting female pheromones, representing the large majority of sensory neurons in their antennae (Leal, 2013). A higher number of sensory neurons is generally assumed to underlie sensitization to the perceived cues (Kudo et al., 2010; Linz et al., 2013; Meisami, 1989; Peichl, 2005). Surprisingly, however, it remains largely untested if and how such neuronal expansions impact sensory processing and behavior.

The drosophilid olfactory system is an excellent model to address these questions (Benton, 2022; Hansson and Stensmyr, 2011). The antenna, the main olfactory organ, houses ∼1000 olfactory sensory neurons (OSNs) that, in *Drosophila melanogaster*, have been classified into ∼50 types based on their expression of one (or occasionally more) Odorant receptors (Ors) or Ionotropic receptors (Irs) (Benton, 2022; Couto et al., 2005; Grabe et al., 2016). The size of individual OSN populations (ranging from ∼10-65 neurons) is stereotyped across individuals, reflecting their genetically hard-wired, developmental programs (Barish and Volkan, 2015; Yan et al., 2020). By contrast, comparisons of homologous OSN types in ecologically-distinct drosophilid species have identified several examples of expansions in OSN populations. Notably, *Drosophila sechellia*, an endemic of the Seychelles that specializes on *Morinda citrifolia* “noni” fruit (Auer et al., 2021; Jones, 2005; Stensmyr, 2009), has an approximately three-fold increase in the neuron populations expressing Or22a, Or85b and Ir75b compared to both *D. melanogaster* and a closer relative, *Drosophila simulans* (Auer et al., 2020; Dekker et al., 2006; Ibba et al., 2010; Prieto-Godino et al., 2017) (**Fig. 1A**). All three of these neuron classes are required for long- and/or short-range odor-guided behaviors (Alvarez-Ocana et al., 2023; Auer et al., 2020) and two of these (Or22a and Ir75b) also display increased sensitivity to noni odors through mutations in the corresponding *D. sechellia* receptor genes (Auer et al., 2020; Dekker et al., 2006; Prieto-Godino et al., 2017). Together, these observations have led to a long-held assumption that these OSN population expansions are important for *D. sechellia*, but this has never been tested.

**Figure 1:**
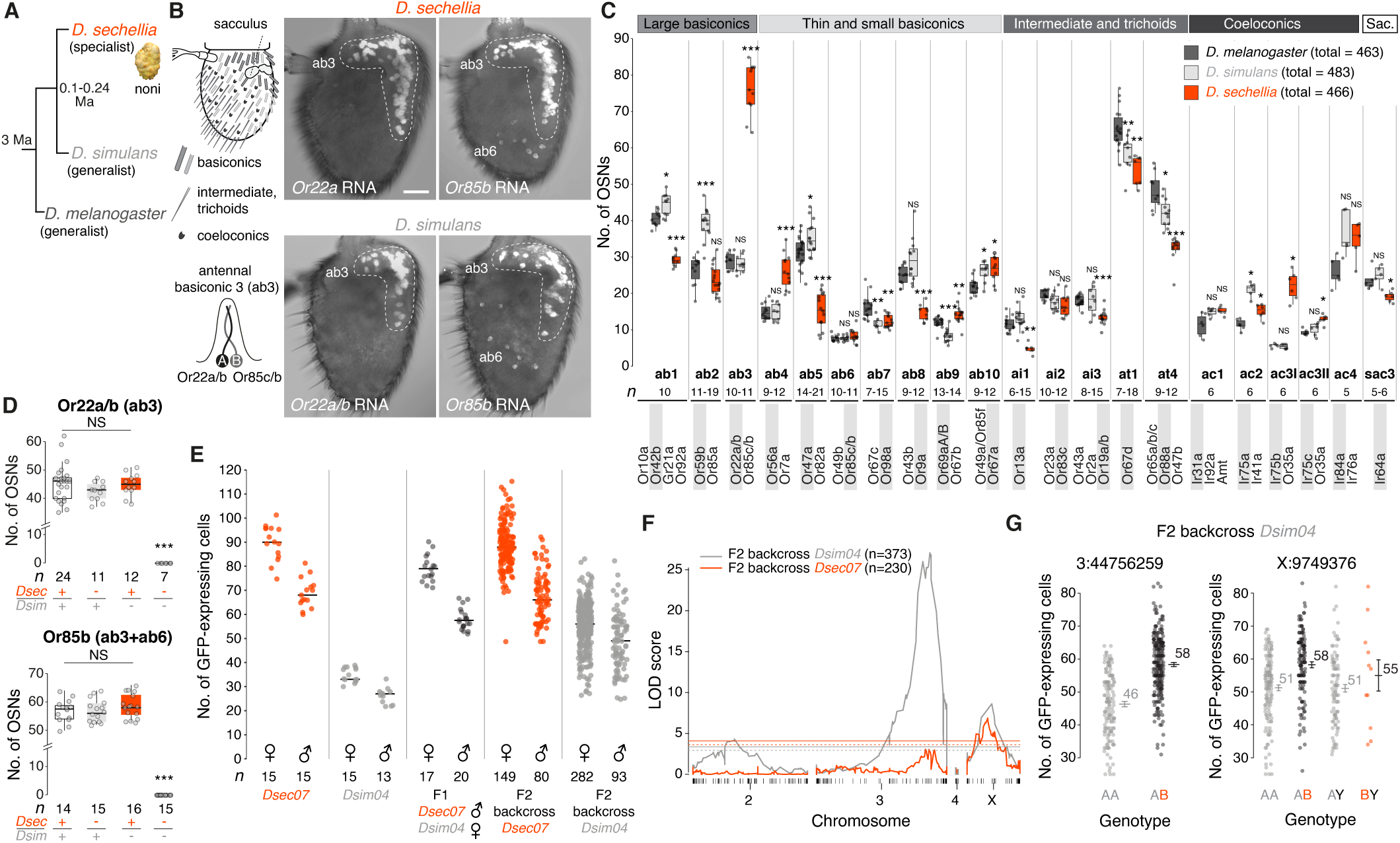
Selective expansion of noni-sensing olfactory sensory neuron populations is a complex developmental trait. **(A)** *D. sechellia* specializes on noni fruit compared to the generalists *D. simulans* and *D. melanogaster*. Ma, million years ago. **(B)** Left top, schematic of the drosophilid third antennal segment covered by sensilla of diverse morphological classes and housing the sacculus. Left bottom, antennal basiconic 3 (ab3) sensilla house two neurons expressing the odorant receptor Or22a (and Or22b in *D. melanogaster* and *D. simulans*; this paralog is lost in *D. sechellia*) in the A neuron (hereafter, “Or22a neuron”) and Or85c/b in the B neuron (both paralogs are co-expressed; hereafter, “Or85b neuron”). Right, antennal *Or22a/(b)* and *Or85b* expression in *D. sechellia* (*Drosophila* Species Stock Center (DSSC) 14021-0248.07, females) and *D. simulans* (DSSC 14021-0251.004, females). Scale bar, 25 µm. In addition to ab3 sensilla (dashed line), Or85b is expressed in ∼10 spatially-segregated OSNs in ab6 sensilla in all three species (**Fig. S1B**). **(C)** Comparison of olfactory sensilla numbers in *D. melanogaster* (*Canton-S*), *D. simulans* (DSSC 14021-0251.004) and *D. sechellia* (DSSC 14021-0248.07) (all females) as assessed by RNA FISH using a diagnostic *Or* probe (grey background) for each sensillum class. Data for Ir-expressing coeloconic sensilla and sacculus chamber 3 (sac3) neurons are from (Prieto-Godino et al., 2017); Ir75d neurons common to ac1, ac2 and ac4 sensilla are not shown. In this and all following panels, box plots show the median and first and third quartiles of the data, overlaid with individual data points. Wilcoxon signed-rank test with comparison to *D. melanogaster*. NS, not significant (*P* > 0.05); \**P* < 0.05; \*\**P* < 0.01; \*\*\**P* < 0.001. **(D)** Reciprocal hemizygosity test of the *Or22a/(b)* and *Or85c/b* loci for contributions to species-specific OSN numbers in *D. simulans*/*D. sechellia* hybrids. Using RNA FISH to quantify numbers of *Or22a/(b)* and *Or85b* expressing OSNs in the indicated genotypes (“*Dsec* +” = *D. sechellia.07* wild-type, “*Dsim* +” = *D. simulans.04* wild-type, “*Dsec* -” = *DsecOr22a^RFP^* or *DsecOr85b^GFP^*, “*Dsim* -” = *DsimOr22a/b^RFP^* or *DsimOr85b^GFP^*), no allele-specific expression differences were observed at either locus. Wilcoxon signed-rank test with comparison to wild-type hybrids. NS, not significant (*P* > 0.05). **(E)** Quantification of GFP-expressing neurons in antennae of *DsecOr85b^GFP^*, *DsimOr85b^GFP^* and F1 hybrid males and females, and in the F2 progeny of the backcrosses of F1 hybrid females to either parental strain. The black line indicates the mean cell number. **(F)** Logarithm of odds (LOD) score across all four chromosomes for loci impacting Or85b neuron numbers based on the phenotypic data in e. Dashed horizontal lines mark *P* = 0.05; non-dashed horizontal lines mark *P* = 0.01. **(G)** Effect sizes for the significant QTL intervals on chromosome 3 and X in the *D. simulans* backcross. A, *D. simulans* allele; B, *D. sechellia* allele. A candidate gene, *lozenge* – encoding a transcription factor involved in sensilla specification development (Gupta et al., 1998) – that is located directly below the highest score of our trait map on the X chromosome, did not affect OSN numbers (Fig. S2E-I).

## Results

### Selective large increases in Or22a and Or85b populations in *D. sechellia*

The increase in Or22a and Or85b OSN numbers in *D. sechellia* reflects the housing of these cells in a common sensory hair, the antennal basiconic 3 (ab3) sensillum (**Fig. 1B, Fig. S1A,B**). To determine how unique this increase is within the antenna, we compared the number of the other ∼20 morphologically-diverse, olfactory sensillar classes – which each house distinct, stereotyped combinations of 1-4 OSN types – in *D. sechellia*, *D. melanogaster* and *D. simulans* through RNA FISH of a diagnostic *Or* per sensillum (combined with published data on *Ir* neurons (Prieto-Godino et al., 2017)) (**Fig. 1C**). We observed several differences in sensillar number between these species, including reductions in ab5, ab8 and ai1 in *D. sechellia*, but only ab3, as well as ac3I that house Ir75b neurons, showed a more than two-fold increase (∼50 more ab3, 2.6-fold increase; ∼15 more ac3I, 3.7-fold increase) in *D. sechellia*. The total number of antennal sensilla is, however, similar across species (**Fig. 1C**).

### OSN population expansion is a complex genetic trait

We next investigated the mechanism underlying the ab3 OSN population expansion in *D. sechellia*. The number of ab3 sensilla was not different when these flies were grown in the presence or absence of noni substrate (**Fig. S1C**), arguing against an environmental influence. We first asked whether the ab3 increase could be explained by simple transformation of sensillar fate. Previous electrophysiological analyses reported loss of ab2 sensilla in *D. sechellia*, which was interpreted as a potential trade-off for the ab3 expansion (Dekker et al., 2006; Stensmyr et al., 2003). However, we readily detected ab2 sensilla histologically in this species (**Fig. 1C, Fig. S1D**), countering this possibility. Moreover, from an antennal developmental fate map in *D. melanogaster* (Chai et al., 2019), we did not observe any obvious spatial relationship between the sensory organ precursors for sensillar classes that display increases or decreases in *D. sechellia* to support a hypothesis of a simple fate switch, although both expanded populations (ab3 and ac3I) originate from peripheral regions of the map (**Fig. S1E**).

We therefore reasoned that genetic changes specifically affecting the development of the ab3 lineage might be involved. We first tested for the existence of species-specific divergence in *cis*-regulation at the receptor loci. Using mutants for both *Or22a/(b)* and *Or85b* in *D. simulans* and *D. sechellia* (Auer et al., 2020), we analyzed receptor expression in interspecific hybrids and reciprocal hemizygotes (lacking transcripts from one or the other receptor allele) (**Fig. 1D, Fig. S1F**). In all hybrid allelic combinations (except those lacking both alleles) we observed a similar number of Or22a and Or85b OSNs, arguing against a substantial contribution of *cis*-regulatory evolution at these loci to the expansion of receptor neuronal expression.

As little is known about the developmental program of ab3 specification, we used an unbiased, whole-genome, quantitative trait locus (QTL) mapping strategy to characterize the genetic basis of the expansion in *D. sechellia*. For high-throughput quantification of ab3 numbers, we generated fluorescent reporters of Or85b neurons in *D. sechellia* and *D. simulans* (**Fig. S2A,B**). Interspecific F1 hybrids displayed an intermediate number of Or85b neurons to the parental strains (**Fig. 1E**). We phenotyped >600 F2 individuals (backcrossed to either *D. sechellia* or *D. simulans*) (**Fig. 1E**), which were then genotyped using multiplexed shotgun sequencing (Andolfatto et al., 2011). The resulting QTL map (**Fig. 1F**) identified two genomic regions linked to variation in Or85b neuron number located on chromosomes 3 and X; these explain a cell number difference of about ∼12 and ∼7 neurons, respectively, between *D. sechellia* and *D. simulans* (**Fig. 1G**). No significant epistasis was detected between these genomic regions (**Methods**) and the relatively low effect size (21.0% and 12.3%, respectively) is consistent with a model in which more than two loci contribute to the species difference in Or85b neuron number. Consistent with the QTL map, introgression of fragments of the *D. sechellia* genomic region spanning the chromosome 3 peak into a *D. simulans* background led to an increased number of Or85b neurons (**Fig. S2C,D**). However, the phenotypic effect was lost with smaller introgressed regions, indicating that multiple loci influencing Or85b neuron number are located within this QTL region (**Fig. S2D**).

In a separate QTL mapping of the genetic basis for the increase in ac3I (Ir75b) neurons in *D. sechellia* (Prieto-Godino et al., 2017) (**Fig. 1C**), we also observed a complex genetic architecture of this interspecific difference, with no evidence for shared loci with the ab3 sensilla expansion (**Fig. S2J,K)**. Thus, in contrast to the evolution of sensory specificity of olfactory pathways – where only one or a few amino acid substitutions in a single receptor can have a phenotypic impact (Alvarez-Ocana et al., 2023; Auer et al., 2020; Prieto-Godino et al., 2017), with some evidence for “hotspot” sites in different receptors (Prieto-Godino et al., 2021) – species differences in the number of OSN types have arisen from distinct evolutionary trajectories involving changes at multiple loci.

### Or22a is required for *D. sechellia* to track host odor

We next investigated if and how increased OSN population size impacts odor-tracking behavior. Previously, we showed that both Or22a and Or85b pathways are essential for long-range attraction to noni in a wind tunnel (Auer et al., 2020). To analyze the behavioral responses of individual animals to odors with greater resolution, we developed a tethered fly assay (Badel et al., 2016) with a timed odor-delivery system (**Fig. 2A**). By measuring the beating amplitude of both wings while presenting a lateralized odor stimulus, we could quantify attractive responses to individual odors, as reflected in an animal’s attempt to turn toward the stimulus source, leading to a positive delta wing beat amplitude (Δ*WBA*) (see **Methods**). We first tested responses of wild-type flies to consecutive short pulses of noni odors, which mimic stimulus dynamics in a plume. As expected, multiple strains of *D. sechellia* displayed attractive responses that persisted through most or all of the series of odor pulses, while *D. melanogaster* strains displayed much more variable degrees of attraction (**Fig. 2B, Fig. S3A**). The persistent attraction in *D. sechellia* was specific to noni odor and could not be observed using apple cider vinegar (**Fig. S3B**).

**Figure 2:**
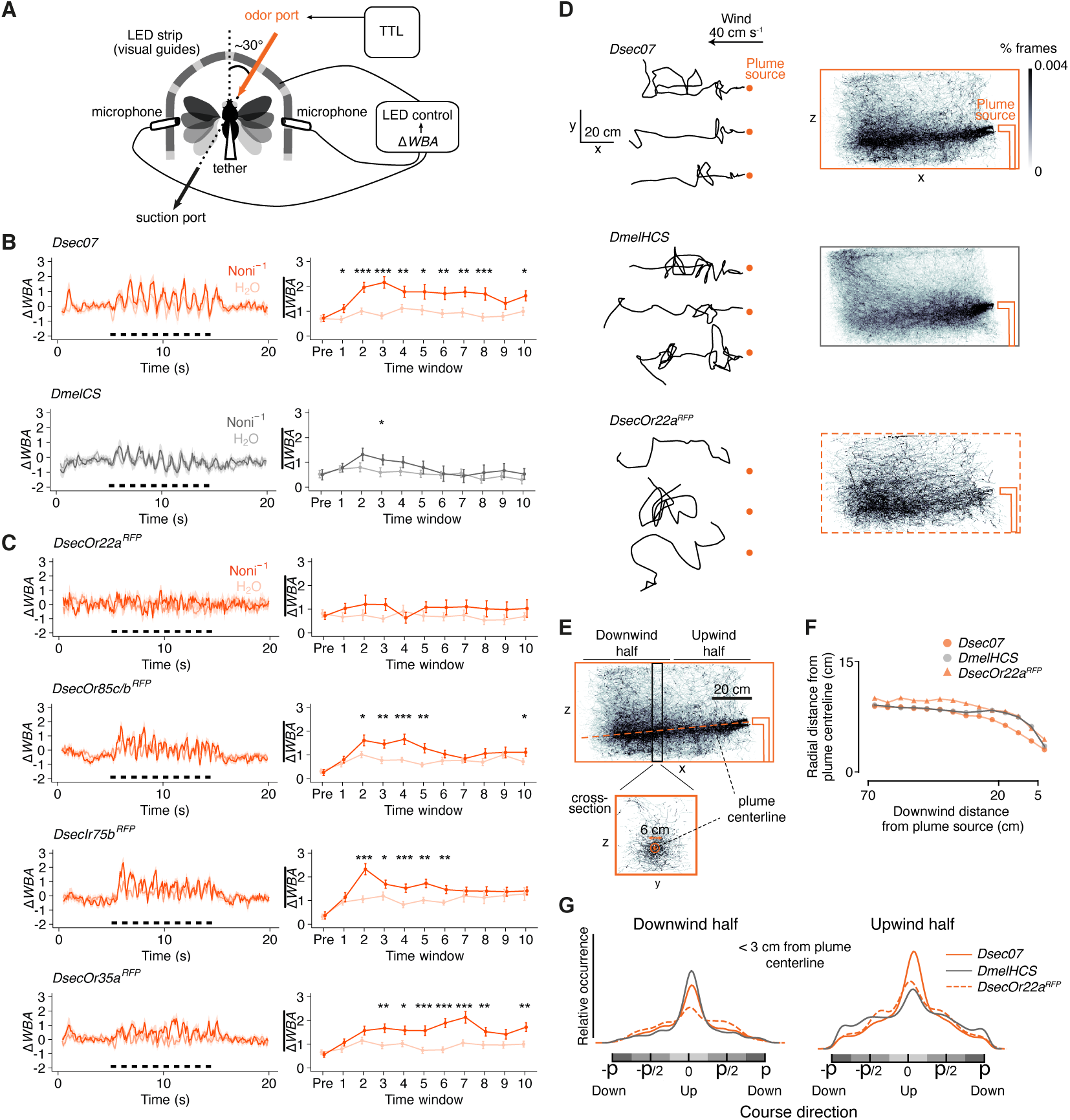
Persistent behavioral tracking of noni odors in *D. sechellia*. **(A)** Schematic of the tethered fly behavioral assay. Δ*WBA*: left-right difference of standardized wing beat amplitudes (see **Methods** for details), TTL: Transistor– transistor logic. **(B)** Odor-tracking behavior towards noni juice and control stimuli (H_2_O) in wild-type *D. sechellia* (DSSC 14021-0248.07) and *D. melanogaster* (CS) flies. Left, time course of Δ*WBA* (mean ± SEM) where black bars indicate the timing of odor stimulation (ten 500 ms pulses with 500 ms intervals). Right, quantification in 1 s time windows immediately prior to stimulus onset (“pre”) and thereafter corresponding to individual stimulus pulses (“1-10”). Mean ± SEM are shown (raw data are provided in the Source Data). Paired *t*-test *** *P* < 0.001; ** *P* < 0.01; * *P* < 0.05, otherwise *P* > 0.05. *n =* 30 animals each. **(C)** Odor-tracking behavior towards noni juice in *D. sechellia Or* and *Ir* mutants. Paired *t*-test, *** *P* < 0.001; ** *P* < 0.01; * *P* < 0.05, otherwise *P* > 0.05. *n* = 30 animals each. **(D)** Left, example trajectories (black lines) in the x-y plane of *D. sechellia* (DSSC 14021-0248.07), *D. melanogaster* (a hybrid *Heisenberg-Canton-S* (*HCS*); our *CS* strain exhibited poor flight performance in this assay) and *D. sechellia Or22a^RFP^* mutants flying in the presence of a noni plume (origin at the orange dot). Right, occupancy heat maps of trajectories that came at least once within 10 cm of the plume centerline for *D. melanogaster* (*n* = 1346 trajectories, 4 recording replicates, 60 flies), *D. sechellia* wild-type (*n* = 835 trajectories, 7 recording replicates, 105 flies) and *D. sechellia Or22a^RFP^* mutants (*n* = 509 trajectories, 6 recording replicates, 90 flies). **(E)** Annotated view of the wild-type *D. sechellia* trajectories (from (D)) illustrating the data analyses performed in (F,G). Based upon the trajectory distribution, we inferred that the noni juice plume sank by ∼5 cm from the odor nozzle to the end of the tracking zone in the wind tunnel; this is likely due to the odor-laden air’s higher water content (and so higher density) than the surrounding air. We therefore estimated the plume center as a line connecting the nozzle and a point 5 cm lower in the z-axis at the end of the wind tunnel. Our results are qualitatively robust whether or not we account for plume sinking (Fig. S4D-F). **(F)** Mean radial distance of the point cloud from the plume centerline for the trajectories in (**D**). Data were binned into 5 cm y-z plane cross-sections starting 5 cm downwind from the plume origin and restricted to 10 cm altitude above or below the estimated plume model. Non-parametric bootstrapped comparison of medians *P* < 0.001. **(G)** Kernel density of the course direction distribution of points within a 3cm radius of the plume centerline (orange circle on cross-section in (E)), further parsed into the point cloud in the downwind or upwind halves of the wind tunnel (see (E)). Downwind half kernels: *D*. *melanogaster* (20,912points from 511 unique trajectories), *D. sechellia* wild-type (15,414 points from 367 trajectories), *D. sechellia Or22a^RFP^* (3,244 points from 164 unique trajectories). Upwind half kernels: *D. melanogaster* (80,594 points from 615 unique trajectories), *D. sechellia* wild-type (136,827 points from 501 unique trajectories), *D. sech^ell^ia Or22a^RFP^* (40,560 points from 286 unique trajectories).

To assess the contribution of distinct olfactory pathways to noni attraction, we tested *D. sechellia* mutants for *Or22a*, *Or85c/b*, *Ir75b* and, as a control, *Or35a*, whose OSN population is also enlarged (as it is paired with Ir75b neurons in ac3I) but is dispensable for noni attraction (Auer et al., 2020). Loss of *Or22a* abolished attraction of flies towards noni, while *Or85c/b* and *Ir75b* mutants retained some, albeit transient, turning towards this stimulus. Flies lacking *Or35a* behaved comparably to wild-type strains (**Fig. 2C**). These results point to Or22a as an important (albeit not the sole) olfactory receptor required for *D. sechellia* to respond behaviorally to noni odor.

To better understand the nature of *D. sechellia* plume-tracking in a more natural setting, we used a wind-tunnel assay combined with 3D animal tracking (Straw et al., 2011) to record trajectories of freely-flying wild-type *D. melanogaster* and *D. sechellia*, as they navigated to the source of a noni juice odor plume (**Fig. 2D, Fig. S4A,B**). Wild-type *D. sechellia* exhibited similar cast and surge dynamics as *D. melanogaster* (van Breugel and Dickinson, 2014) (**Fig. 2D**). However, we observed that *D. sechellia* maintained flight paths closer to the plume centerline (**Fig. 2D-F**), a difference that was particularly evident close (<20 cm) to the plume source (**Fig. 2D-F**). Consistent with the phenotype observed in the tethered-fly assay, *D. sechellia Or22a* mutants lacked strong plume-tracking responses (**Fig. 2D-F**), while not exhibiting obvious impairment in flight performance (**Fig. S4C**). To extend analysis of these observations we quantified the distribution of flies within a 3 cm radius of the estimated odor plume centerline in the downwind and upwind halves of the wind tunnel (**Fig. 2E,G**). In the downwind half, wild-type *D. sechellia* and *D. melanogaster* had comparable course direction distributions (**Fig. 2G**). By contrast, in the upwind half, *D. sechellia* maintained a tighter upwind course distribution suggesting they were more likely to be in an upwind surging state compared to *D. melanogaster* as they approach the odorant source (**Fig. 2G**). *D. sechellia Or22a* mutants did not appear to be strongly oriented into the wind, notably in the downwind half, further indicating the importance of this olfactory pathway in stereotypical plume-tracking behaviors (**Fig. 2G**).

### *D. sechellia*’s increase in OSN number is important for persistent odor-tracking behavior

To investigate the importance of Or22a neuron number for odor-tracking behavior we expressed CsChrimson in Or22a neurons in *D. melanogaster* and *D. sechellia* to enable specific stimulation of this pathway with red light. This optogenetic approach had two advantages: first, it allowed us to calibrate light intensity to ensure equivalent Or22a OSN activation between species; second, because only Or22a neurons are activated by light, we could eliminate sensory contributions from other olfactory pathways that have overlapping odor tuning profiles to Or22a.

We first confirmed light-evoked spiking in Or22a neurons in *D. melanogaster* and *D. sechellia* and determined the light intensity evoking equivalent spike rates (**Fig. 3A**). We then performed lateralized optogenetic activation (Gaudry et al., 2013) of Or22a OSNs in the tethered fly assay, mimicking lateralized odor input by focusing the light beam on one antenna (**Fig. 3A, Fig. S5A**). Pulsed optogenetic activation of *D. sechellia* Or22a OSNs induced attractive behavior with a similar magnitude and dynamics as pulsed odor stimuli (**Fig. 3B**), demonstrating the sufficiency of this single olfactory pathway for evoking behavior. Notably, the attractive behavior was more persistent over the series of light pulses in *D. sechellia* than in *D. melanogaster* (**Fig. 3A, Fig. S5B-D**), consistent with a hypothesis that a higher number of Or22a OSNs supports enduring behavioral attraction to a repeated stimulus.

**Figure 3:**
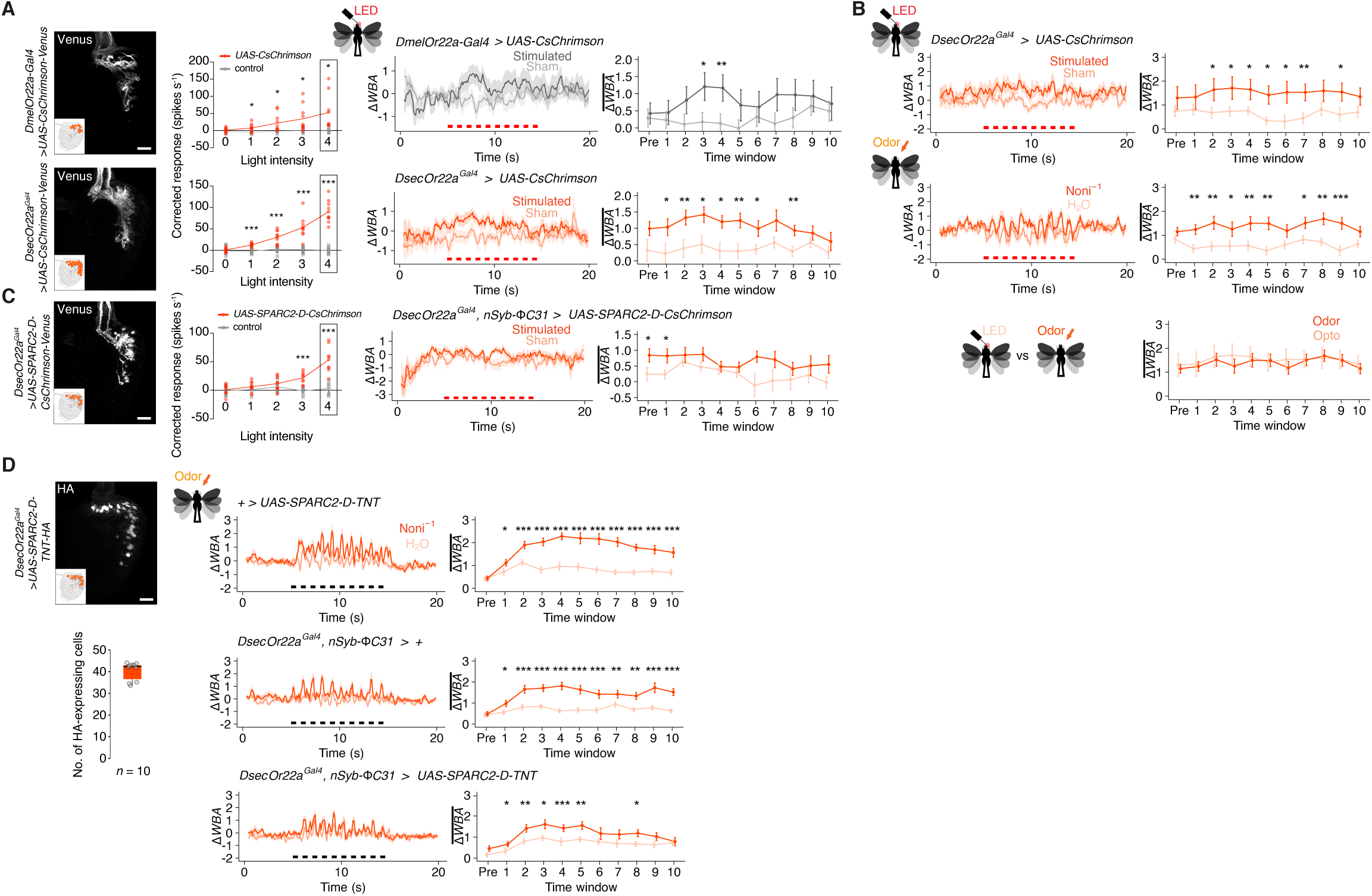
Behavioral significance of OSN number. **(A)** Optogenetic stimulation of Or22a OSNs in *D. melanogaster* (top) and *D. sechellia* (bottom). Left, expression of CsChrimson in the antenna detected by expression of the Venus tag. Scale bar, 20 µm. Middle, single-sensillum recordings of Or22a OSNs in response to optogenetic stimulation of CsChrimson-expressing and control sensilla. The red line links the mean neuronal response at each light intensity, overlaid with individual data points. The black frame indicates the light intensity used for behavioral experiments. *n* = 5-10 sensilla (exact numbers are listed in **Supplemental Table 1**), unpaired Student’s *t*-test, *** *P* < 0.001; * *P* < 0.05, otherwise *P* > 0.05. Right, behavioral responses of the same genotypes in response to red light stimulation (ten 500 ms pulses with 500 ms intervals, indicated by the red bars). Paired *t*-test, ** *P* < 0.01; * *P* < 0.05, otherwise *P* > 0.05. *n =* 29 (*D. melanogaster*) and 30 (*D. sechellia*) animals. Genotypes: *D. melanogaster w;UAS-CsChrimson-Venus/+* (control), *w;UAS-CsChrimson-Venus/Or22a-Gal4* (experimental), *D. sechellia w;;UAS-CsChrimson-Venus/+* (control), *w;Or22a^Gal4^/+;UAS-CsChrimson-Venus/+* (experimental). **(B)** Behavioral responses of *D. sechellia* upon optogenetic stimulation of Or22a OSNs (top) and to noni odor stimulation (middle), in the same animals. Bottom, comparison of Δ*WBA* between light and odor responses. Paired *t*-test, *** *P* < 0.001, ** *P* < 0.01, * *P* < 0.05, otherwise *P* > 0.05. *n* = 27 animals each. Genotype as in (**A**). **(C)** Sparse activation of *D. sechellia* Or22a OSNs. Left, expression of CsChrimson in an antenna of *D. sechellia* expressing *UAS-SPARC2-D-CsChrimson-Venus* in Or22a OSNs. Fluorescent labelling was sparser than in *UAS-CsChrimson-Venus* expressing animals, but the dense packing of membrane-labelled neurons prevented quantification. Experiments in *D. melanogaster* (**Fig. S6B**) and with the *UAS-SPARC2-D-TNT-HA* transgene (below) support expression in ∼50% of Or22a OSNs with this SPARC version. Scale bar, 20 µm. Middle, single-sensillum recordings of Or22a OSNs in response to optogenetic stimulation. ab3 sensilla were first identified by stimulation with diagnostic odors (not shown); responses of CsChrimson-expressing neurons (experimental group) and non-expressing neurons (control, often from the same animal) are shown. *n* = 8-9 (exact numbers are listed in **Supplemental Table 1**), unpaired *t*-test, *** *P* < 0.001; * *P* < 0.05, otherwise *P* > 0.05. Right, *D. sechellia* behavior upon optogenetic activation of about half of their Or22a expressing neurons. Paired *t*-test, *** *P* < 0.001; * *P* < 0.05; otherwise *P* > 0.05. *n =* 26 animals. Genotypes: *D. sechellia w;Or22a^Gal4^/UAS-SPARC2-D-CsChrimson-Venus;;nSyb-*Φ*C31/+*. **(D)** Sparse inhibition of *D. sechellia* Or22a OSNs. Left, HA immunofluorescence in an antenna of *D. sechellia* (females) expressing *UAS-SPARC2-D-TNT-HA* in Or22a OSNs. Scale bar, 20 µm. Quantification of cell-labelling is shown below. Middle, *D. sechellia* odor-tracking behavior towards noni juice of flies in effector control (top), driver control (middle), or in experimental animals with blocked synaptic transmission in approximately half of their Or22a neurons (bottom). Paired *t*-test, *** *P* < 0.001; ** *P* < 0.01; * *P* < 0.05; otherwise *P* > 0.05. *n =* 34 animals each. Genotypes: *D. sechellia w;UAS-SPARC2-D-TNT-HA-GeCO/+;;+/+* (effector control), *D. sechellia w;Or22a^Gal4^/+;;nSyb-*Φ*C31/+* (driver control), *D. sechellia w;Or22a^Gal4^/UAS-SPARC2-D-TNT-HA-GeCO;;nSyb-*Φ*C31/+* (experimental group).

To explicitly test this possibility, we generated genetic tools to reproducibly manipulate the number of active Or22a OSNs in *D. sechellia*, using the SPARC technology developed in *D. melanogaster* (Isaacman-Beck et al., 2020) (**Fig. S6A**). Combining a *SPARC2-D-CsChrimson* transgene with *Or22a-Gal4* allowed us to optogenetically stimulate ∼50% of these neurons (**Fig. 3C, Fig. S6B**). Although we confirmed light-evoked Or22a neuron spiking in these animals, they did not display attractive behavior towards the light stimulus, in contrast to similar stimulation of all Or22a OSNs (**Fig. 3C, Fig. S6C**). This result implies that the number of active OSNs is critical to induce attraction in *D. sechellia*.

Optogenetic activation of Or22a neurons only partially mimics differences between species because it does not offer the opportunity for any possible plasticity in circuit properties that are commensurate with differences in OSN number (as described below). We therefore took a complementary approach, through neuronal silencing, using a *SPARC2-D-Tetanus Toxin* (*TNT*) transgene. With this tool we could silence on average ∼50% of Or22a OSNs (**Fig. 3D**) – likely from mid/late-pupal development (when the *Or22a* promoter is activated (Pan et al., 2017)) – without directly inhibiting other peripheral or central neurons (**Fig. S6D**). Importantly, when tested in our tethered flight assay, these flies showed weaker and more transient attraction towards noni odor, contrasting with the persistent noni attraction of control animals (**Fig. 3D**). Together these results support the hypothesis that the increased Or22a OSN number observed in *D. sechellia* is essential for strong and sustained attractive olfactory behavior.

### OSN population expansions lead to increased pooling on cognate projection neurons

To better understand how increased OSN number might enhance odor-guided behavior, we first characterized the anatomical properties of the circuitry. The axons of OSNs expressing the same receptor converge onto a common glomerulus within the antennal lobe in the brain (Schlegel et al., 2021) (**Fig. 4A**). Here, they form cholinergic synapses on mostly uniglomerular projection neurons (PNs) – which transmit sensory signals to higher olfactory centers – as well as on broadly-innervating local interneurons (LNs) and a small proportion on other OSNs (Mosca and Luo, 2014; Schlegel et al., 2021). To visualize PNs in *D. sechellia*, we generated specific driver transgenes in this species using constructs previously-characterized in *D. melanogaster* (Elkahlah et al., 2020; Tirian and Dickson, 2017): *VT033006-Gal4*, which drives expression in many uniglomerular PN types and *VT033008-Gal4* which has sparser PN expression (**Fig. 4B**). Using these drivers to express photoactivable-GFP in PNs, we performed spatially-limited photoactivation of the DM2 and VM5d glomeruli – which receive input from Or22a and Or85b OSNs, respectively (**Fig. 4A**) – to visualize the partner PNs. We confirmed that DM2 is innervated by 2 PNs in both *D. melanogaster* and *D. sechellia* (Auer et al., 2020) and further found that VM5d also has a conserved number of PNs (on average 4) in these species (**Fig. 4C**). Together with data that the Ir75b glomerulus, DL2d, has the same number of PNs in *D. melanogaster*, *D. simulans* and *D. sechellia* (Ellis et al., 2023; Prieto-Godino et al., 2017), these observations indicate that *D. sechellia*’s OSN population expansions are not accompanied by increases in PN partners.

**Figure 4:**
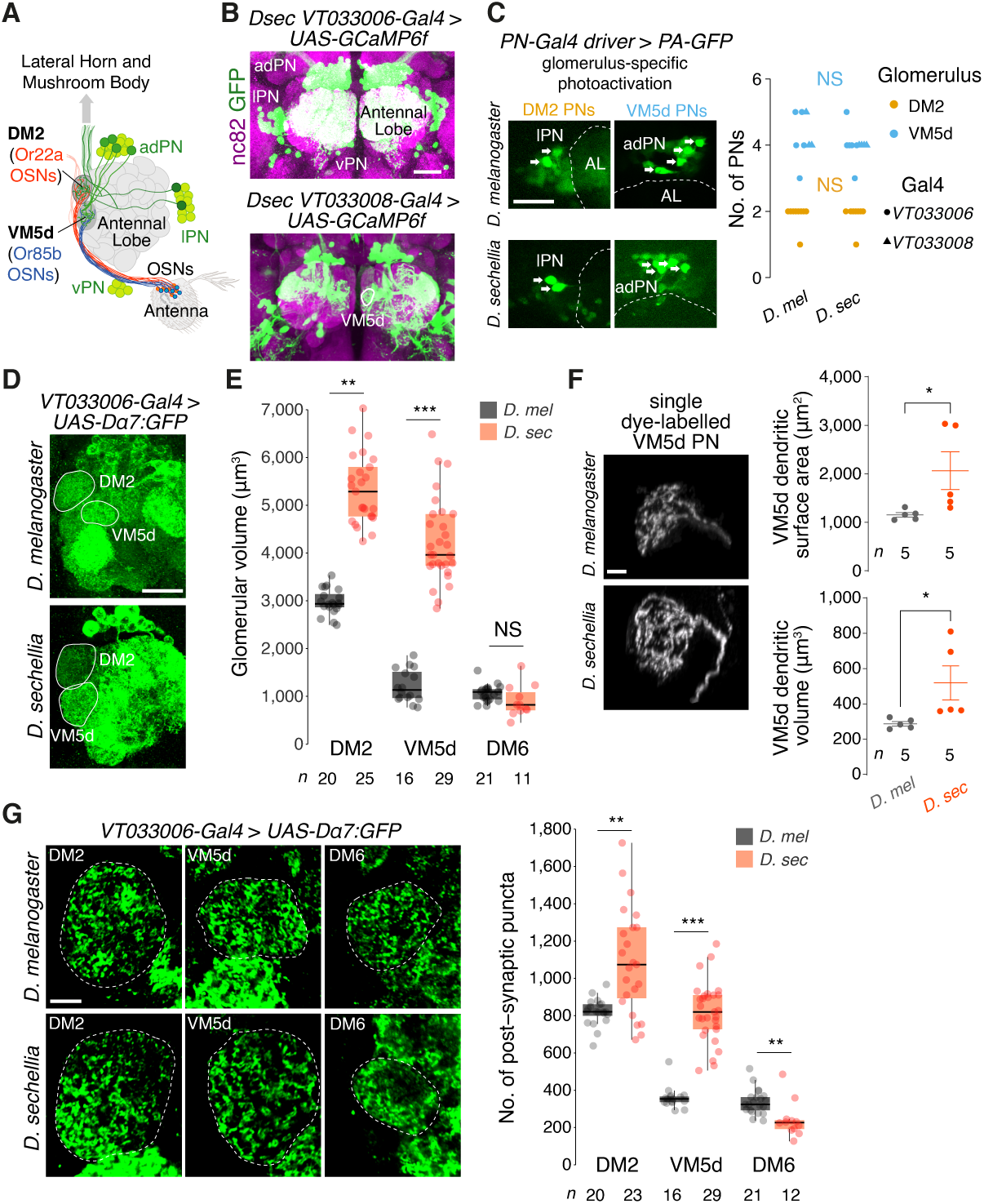
Increased sensory and synaptic pooling in noni-sensing glomeruli. **(A)** Schematic of OSN-PN connectivity in the antennal lobe. Or22a OSNs (orange) and Or85b OSNs (blue) have cell bodies in the antenna and axons projecting to the DM2 and VM5d glomerulus, respectively. The soma of second-order PNs (green) are located in three distinct clusters around the antennal lobe (ad, anterodorsal; l, lateral; v, ventral); these neurons synapse with OSNs (the majority constrained to a single glomerulus) and send axonal projections to higher olfactory centers. For clarity, LN are not illustrated (see text) **(B)** Transgenic labelling of PNs in the *D. sechellia* antennal lobe using the *VT033006-Gal4* or *VT033008-Gal4* drivers to express *UAS-GCaMP6f* (here used simply as a fluorescent reporter). Immunofluorescence on whole-mount brains was performed with antibodies against GFP (detecting GCaMP6f) and nc82 (labelling the synaptic protein Bruchpilot (Wagh et al., 2006)). The VM5d glomerulus is demarcated by a white line in the *VT033008-Gal4* line. Scale bar, 25 µm. **(C)** Left, representative image of VM5d PNs labelled by photo-activatable GFP (PA-GFP) in *D. melanogaster* and *D. sechellia*. Genotypes: *D. melanogaster UAS-C3PA-GFP/+;UAS-C3PA-GFP/VT033008-Gal4* (for VM5d PNs) or *UAS-C3PA-GFP/+;UAS-C3PA-GFP/VT033006-Gal4* (for DM2 PNs); *D. sechellia UAS-C3PA-GFP/VT033008-Gal4* (for VM5d PNs) or *UAS-C3PA-GFP/VT033006-Gal4* (for DM2 PNs). Arrows indicate the PN cell bodies; faint background GFP signal in other soma are irrelevant neuron types. The antennal lobe (AL) is demarcated by a dashed line. Scale bar, 25 µm. Right, quantification of Or22a and Or85c/b PN numbers innervating the DM2 and VM5d glomeruli, respectively, by labelling with PA-GFP (using two different driver lines for VM5d PNs) in *D. sechellia* and *D. melanogaster*. Mann-Whitney *U*-test, NS, not significant. *P* > 0.05. **(D)** Visualization of antennal lobe glomeruli by expression of the Dα7-GFP post-synaptic marker in PNs. Genotypes: *D. melanogaster w;;VT033006-Gal4/UAS-Dα7-GFP*; *D. sechellia w;VT033006-Gal4/+;UAS-Dα7-GFP*/+. Scale bar, 20 µm. **(E)** Quantification of the volumes of DM2, VM5d and a control glomerulus, DM6 (innervated by Or67a neurons) in *D. sechellia* and *D. melanogaster* (females). Wilcoxon signed-rank test. \*\**P* < 0.005; \*\*\**P* < 0.001. **(F)** Representative images of single dye-labelled VM5d PNs in *D. melanogaster* and *D. sechellia* Genotypes: *D. melanogaster w;VM5d-Gal4/UAS-GFP*, *D. sechellia w;VM5d-Gal4/UAS-myrGFP* (the GFP fluorescence is not shown). Scale bar, 5 µm. Right, quantification of VM5d PN dendrite surface area and volume. Student’s *t*-test. \**P* < 0.05. **(G)** Left, representative images of post-synaptic puncta in VM5d, DM2 and DM6 PNs labelled by Dα7-GFP in *D. melanogaster* and *D. sechellia* (genotypes as in (**D**)). Scale bar, 5 µm. Right, quantification of the number of post-synaptic puncta in these glomeruli. Wilcoxon signed-rank test. \*\**P* < 0.005; \*\*\**P* < 0.001.

Next, we expressed a post-synaptic marker, the GFP-tagged Dα7 acetylcholine receptor subunit, in PNs (Leiss et al., 2009; Mosca et al., 2017; Mosca and Luo, 2014) in *D. melanogaster* and *D. sechellia* (**Fig. 4D**). Quantification of glomerular volume as visualized with this reporter confirmed previous observations, using OSN markers (Auer et al., 2020; Dekker et al., 2006; Ellis et al., 2023; Ibba et al., 2010; Prieto-Godino et al., 2017), that the DM2 and VM5d glomeruli, but not a control glomerulus (DM6), are specifically increased in *D. sechellia* compared to *D. melanogaster* (**Fig. 4E**). Given the constancy in PN number (**Fig. 4C**), this observation implied that PN dendrites must exhibit anatomical differences to occupy a larger volume. We examined this possibility through visualization of single VM5d PNs by dye-labelling (in the course of electrophysiology experiments described below). Reconstruction of dendrite morphologies revealed that *D. sechellia* VM5d PNs have increased dendritic surface area and volume compared to the homologous neurons in *D. melanogaster* (**Fig. 4F**).

Finally, we quantified post-synaptic puncta of Dα7:GFP to estimate the number of excitatory OSN-PN connections in these glomeruli. Both DM2 and VM5d, but not DM6, displayed more puncta in *D. sechellia* than in *D. melanogaster* (**Fig. 4G**). Although quantifications of fluorescent puncta are likely to substantially underestimate the number of synapses detectable by electron microscopy (Mosca and Luo, 2014; Schlegel et al., 2021), this reporter should still reflect the relative difference between species. Together these data suggest that an increase in OSN numbers leads to larger glomerular volumes, increased dendritic arborization in partner PNs and overall more synaptic connections between OSNs and PNs.

### OSN number increases do not lead to sensitization of PN responses

To investigate the physiological significance of these OSN-PN circuit changes, we generated genetic reagents to visualize and thereby perform targeted electrophysiological recordings from specific PNs. We focused on VM5d PNs due to the availability of an enhancer-Gal4 transgene for sparse genetic labelling of this cell type (Li et al., 2020). Moreover, the partner Or85b OSNs’ sensitivities to the best-known agonist, *2*-heptanone, are indistinguishable between species (Auer et al., 2020), enabling specific assessment of the impact of OSN population expansion on PN responses. Through whole-cell patch clamp recordings from VM5d PNs in *D. melanogaster* and *D. sechellia* (**Fig. 5A**), we first observed that the input resistance of these cells was ∼2-fold lower in *D. sechellia* (**Fig. 5B**), consistent with their larger dendritic surface area and volume (**Fig. 4F**). The resting membrane potential (**Fig. 5C**) and spontaneous activity (**Fig. 5D**) of these PNs were, however, unchanged between species. Surprisingly, VM5d PNs displayed no obvious increase in odor sensitivity in *D. sechellia* (**Fig. 5E, Fig. S7A**); if anything, the peak spiking rate (within the first 50 ms after PN response onset) tended to be higher in *D. melanogaster*. However, we observed that the PN firing during the odor stimulus displayed a starker decay in *D. melanogaster* than *D. sechellia* (**Fig. 5F, Fig. S7B**). These data support a model in which increased synaptic input by more OSNs in *D. sechellia* is compensated by decreased PN input resistance precluding the sensitization of responses in this cell type. Instead, OSN increases might impact the temporal dynamics of PN responses (explored further below).

**Figure 5:**
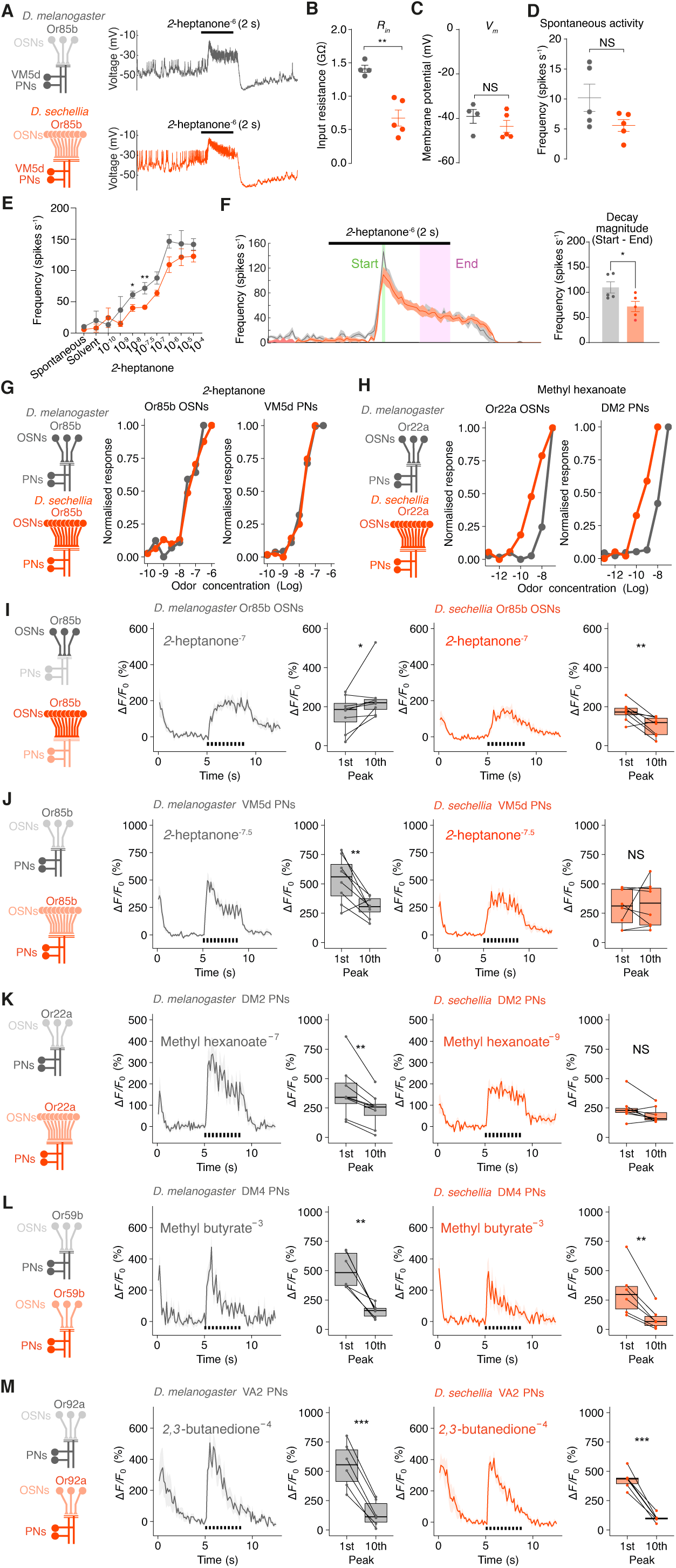
Sustained representation of noni odor stimuli in PNs of *D. sechellia*. **(A-D)** Whole-cell patch clamp recording from VM5d PNs in *D. melanogaster* and *D. sechellia*; the glomerular circuit is schematized on the left. Genotypes as in **Fig. 4F**. (**A**) Voltage trace of VM5d PNs in response to a pulse of *2*-heptanone. (**B-D**), Comparison of input resistance (**B**), resting membrane potential (**C**), and spontaneous activity (**D**) between *D. melanogaster* and *D. sechellia*. Student’s *t*-test. \*\**P* < 0.01. *n* = 4-5. **(E)** Dose-response relationship of VM5dPN firing to *2*-heptanone. Quantification of spike frequency was performed in a 50 ms window covering the peak response. Student’s *t*-test. \*\**P* < 0.01, \**P* < 0.05. *n* =2-5 animals (exact numbers and mean responses are listed in Supplemental Table 1). EC_50_ values [Log] are as follows: -7.46 (*Dmel*), -6.87 (*Dsec*). **(F)** VM5d PN spike frequency in response to 10^-6^ dilution of *2*-heptanone. Left, time course of spike frequency. Mean ± SEM are shown. Right, quantification of the decay magnitude (i.e. start (first 50 ms) - end (last 500 ms before odor offset)). Student’s *t*-test. \**P* < 0.05. *n* = 5 animals each. **(G-H)** Dose-dependent, odor-evoked calcium responses in Or85b/VM5d (**G**) and Or22a/DM2 (**H**) OSN axon termini and PN dendrites in the antennal lobe of *D. melanogaster* and *D. sechellia*, reported as normalized GCaMP6f fluorescence changes. Plots are based on the data in **Fig. S8**. EC_50_ values [Log] are as follows: Or85b OSNs: -6.50 (*Dmel*), -6.51 (*Dsec*); VM5d PNs: -7.82 (*Dmel*), -7.87 (*Dsec*); Or22a OSNs: -7.42 (*Dmel*), -8.23 (*Dsec*); DM2 PNs: -7.97 (*Dmel*), -9.61 (*Dsec*). Genotypes are as follows. OSN calcium imaging: *D. melanogaster UAS-GCaMP6f/Orco-Gal4*,*UAS-GCaMP6f*, *D. sechellia UAS-GCaMP6f/UAS-GCaMP6f;;+/DsecOrco^Gal4^*. PN calcium imaging: *D. melanogaster UAS-GCaMP6f/+;+/VT033008-Gal4* (for VM5d PNs), *D. melanogaster UAS-GCaMP6f/+;+/VT033006-Gal4* (for DM2 PNs), *D. sechellia UAS-GCaMP6f/VT033008-Gal4* (for VM5d PNs), *UAS-GCaMP6f/VT033006-Gal4* (for DM2 PNs). **(I)** Responses of Or85b OSNs to pulsed odor stimuli (ten 200 ms pulses, each separated by 200 ms, as indicated by the black bars). For both species, left panels show the time course (mean ± SEM Δ*F/F_0_*) and right panels show the quantification of Δ*F/F_0_* peak to the 1^st^ and 10^th^ stimulation. Paired *t*-test. ** *P* < 0.01, * *P* < 0.05. *n* = 8 animals each. **(J-K)** Pulsed odor responses of VM5d PNs (J) and DM2 PNs (K). Paired *t*-test, ** *P* < 0.01. *n =* 8 animals each. **(L-M)** Pulsed odor responses of VA2 (Or92a) and DM4 (Or59b) PNs (two control pathways where the number of cognate OSNs is conserved between species (**Fig. 1C**)). Paired *t*-test. *** *P* < 0.001; ** *P* < 0.01. *n =* 6 animals each. Dose response data for these neurons are provided in **Fig. S11**. Genotypes are as for DM2 PN imaging in (**H**).

To substantiate and extend these observations, we expressed GCaMP6f broadly in OSNs or PNs and measured odor-evoked responses in specific glomeruli using two-photon calcium imaging. When measuring calcium responses to a short pulse of *2*-heptanone in Or85b OSN axon termini in the VM5d glomerulus, we observed, as expected (Auer et al., 2020), no sensitivity differences (**Fig. 5G, Fig. S8A**). VM5d PNs also displayed no obvious increase in odor sensitivity in *D. sechellia* (**Fig. 5G, Fig. S8B**), consistent with our patch clamp recordings (**Fig. 5E**). Next, we investigated Or22a OSNs and DM2 PNs after stimulation with the noni odor methyl hexanoate (Dekker et al., 2006). In this olfactory pathway, *D. sechellia* OSNs displayed responses at odor concentrations approximately two orders of magnitude lower than in *D. melanogaster* (**Fig. 5H, Fig. S8C**), concordant with previous electrophysiological analyses (Auer et al., 2020; Dekker et al., 2006; Stensmyr et al., 2003). *D. sechellia* DM2 PNs displayed a similar degree of heightened sensitivity compared to those in *D. melanogaster*, supporting that an increased Or22a OSN number does not lead to further sensitization of these PNs (**Fig. 5H, Fig. S8D**). To test the sufficiency of differences in receptor properties (Auer et al., 2020) to explain PN activity differences, we expressed *D. melanogaster* or *D. sechellia* Or22a in *D. melanogaster* Or22a neurons lacking the endogenous receptors. This manipulation conferred a species-specific odor response profile to these OSNs (Auer et al., 2020). Measurement of responses to methyl hexanoate in DM2 PNs revealed higher sensitivity in animals expressing *D. sechellia* Or22a compared to those expressing *D. melanogaster* Or22a (**Fig. S8E**); notably, this difference was similar in magnitude to the endogenous sensitivity differences of *D. sechellia* and *D. melanogaster* DM2 PNs. Together, the analyses of Or85b and Or22a pathways indicate that a larger OSN population does not contribute to enhanced PN sensitization.

### OSN number increases do not lead to increased reliability of PN responses

Pooling of inputs on interneurons has also been suggested to reduce the noise in central sensory representations, since spontaneous activity of each OSN is uncorrelated and becomes averaged out as OSN activities are summated at PNs (Bhandawat et al., 2007; Jeanne and Wilson, 2015; Serences, 2011). This phenomenon should reduce the variability of PN response magnitude across multiple odor presentations. We therefore examined whether increased OSN number plays a role in reducing trial-to-trial variability in PN responses by comparing odor responses, and their variation, in VM5d PNs to eight trials of *2-* heptanone stimulation. These experiments did not reveal any differences in PN response reliability between *D. melanogaster* and *D. sechellia* (**Fig. S9**). Consistent with this calcium imaging analyses, the lack of a significant difference in VM5d PN spontaneous spiking frequency between species (**Fig. 5D**) also argues that increased sensory pooling in *D. sechellia* does not substantially reduce noise in this circuitry.

### Pathways with increased OSN number display reduced decay magnitude of PN responses

Most natural odors exist as turbulent plumes, which stimulate OSNs with complex, pulsatile temporal dynamics (van Breugel and Dickinson, 2014). To test if odor temporal dynamics in these olfactory pathways are influenced by OSN number, we repeated the calcium imaging experiments using ten consecutive, short pulses of odor. Or85b OSNs displayed a plateaued response to pulses of *2*-heptanone in both *D. melanogaster* and *D. sechellia*, albeit with a slight decay over time in the latter species (**Fig. 5I**). *D. melanogaster* VM5d PNs showed decreasing responses following repeated exposure to short pulses, presumably reflecting adaptation, as observed in multiple PN types (Kazama and Wilson, 2008). By contrast, VM5d PNs in *D. sechellia* displayed responses of similar magnitude throughout the series of odor pulses (**Fig. 5J**). This species difference in PN responses was also seen with long-lasting odor stimulation (**Fig. S10**) and is consistent with a smaller difference between spiking frequencies at the start and end of odor stimulation in *D. sechellia* VM5d PNs measured by electrophysiological recordings (**Fig. 5F and S7B**). Imaging the responses of Or22a OSN partner PNs in DM2 to pulsed odor stimuli (using concentrations of methyl hexanoate that evoked similar activity levels between the species (**Fig. 5H**)) revealed a similar result: *D. melanogaster* DM2 PN responses decreased over time while *D. sechellia* DM2 PNs responses were unchanged in magnitude (**Fig. 5K**). However, for two olfactory pathways where the numbers of cognate OSNs are not increased in *D. sechellia* (Or59b (DM4) and Or92a (VA2) (**Fig. 1C**)), the PNs in both species displayed decreasing responses over the course of stimulations (**Fig. 5L,M, Fig. S11**). Together, our data indicate that OSN number increases in *D. sechellia*’s noni-sensing pathways might result, directly or indirectly, in reduced decay magnitude of PN responses to dynamic or long-lasting odor stimuli.

### Species-specific PN response properties are due, at least in part, to differences in lateral inhibition

More sustained PN responses in *D. sechellia* could be due to species differences in several aspects of glomerular processing, involving OSNs, PNs and/or LNs. We examined this phenomenon through calcium imaging in VM5d PNs before and after pharmacological inhibition of different synaptic components. Blockage of inhibitory neurotransmission – which is principally mediated by LNs acting broadly across glomeruli (Liu and Wilson, 2013; Olsen et al., 2010; Wilson and Laurent, 2005) – decreased adaptation in *D. melanogaster* VM5d PNs, as expected. This effect was predominantly due to inhibition of GABA_B_ rather than GABA_A_/GluCl receptors (**Fig. S12A,B**). By contrast, inhibitory neurotransmitter receptor blockage did not lead to changes in temporal dynamics of responses in *D. sechellia* (**Fig. 6A and Fig. S12A,B**). These observations suggest that differences in the strength of inhibition between species contribute to differences in PN decay magnitude (see Discussion). We note that *D. sechellia* PNs displayed apparent decreases in odor response magnitude (as measured by *ΔF/F*) upon pharmacological treatment (**Fig. 6A**), but this effect is most likely due to elevated baseline activity (*F_0_*) in this species (**Fig. S12C**).

**Figure 6:**
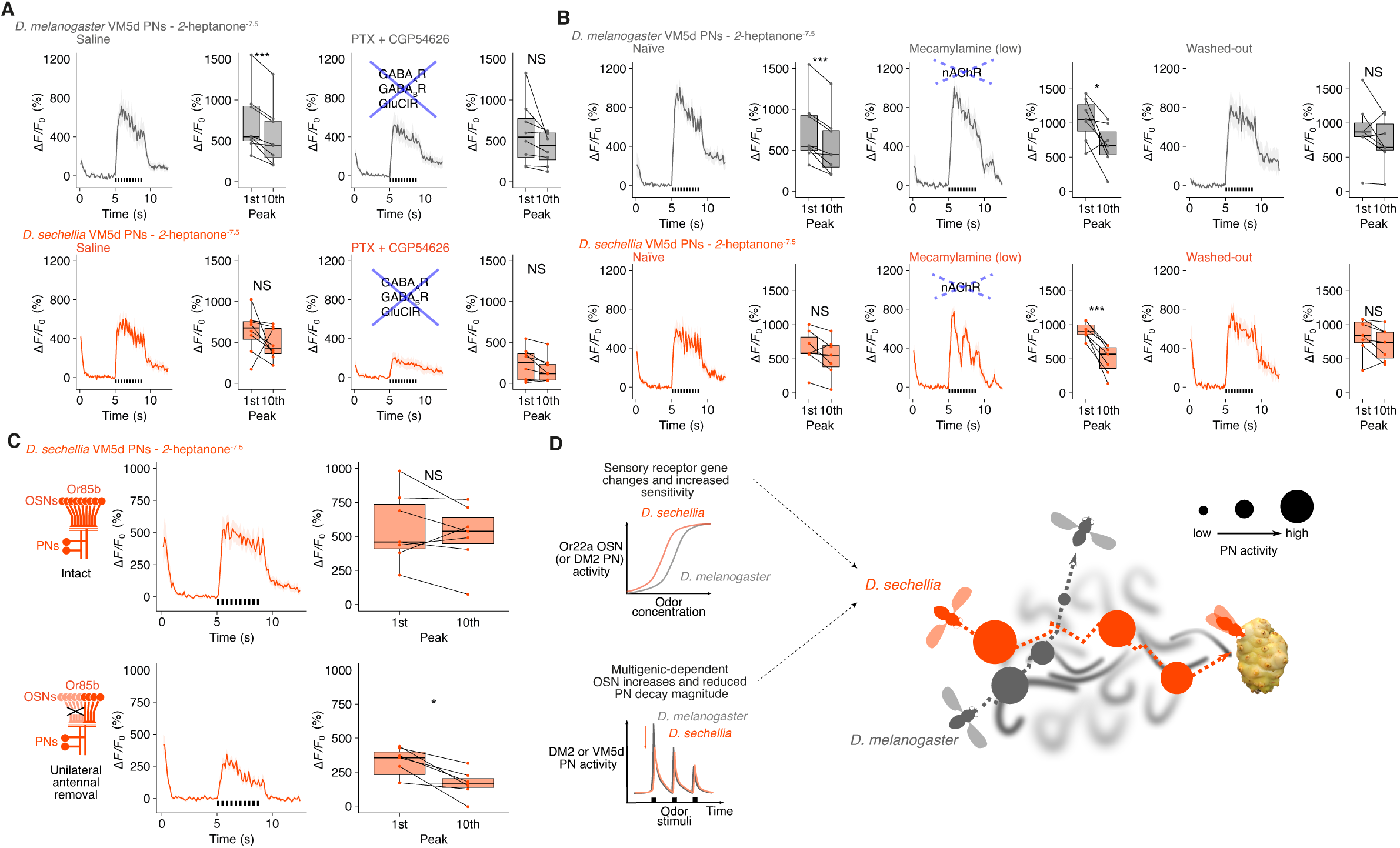
Mechanisms of sustained PN responses in *D. sechellia*. **(A)** Odor pulse responses of VM5d PNs following application of a GABA antagonist. PN responses in normal AHL saline (left) or containing 100 µM picrotoxin + 50 µM CGP54626 (right). Paired *t*-test, *** *P* < 0.001, NS *P* > 0.05. *n =* 8 animals each. Genotypes are indicated in **Fig. 5G** legend. **(B)** Odor pulse responses of VM5d PNs following application of low doses (200 µM) of mecamylamine (nAChR antagonist) to weakly block cholinergic inputs. *D. melanogaster* and *D. sechellia* PN responses in normal AHL saline, mecamylamine and AHL saline wash-out. Paired *t*-test. *** *P* < 0.001, * *P* < 0.05, NS *P* > 0.05. *n =* 7 animals each. **(C)** Odor pulse responses of *D. sechellia* VM5d PNs in intact (top) and right antenna-ablated (bottom) animals, in which the OSN input is halved. Paired *t*-test, * *P* < 0.05, NS *P* > 0.05. *n =* 7 each. **(D)** Model illustrating the complementary effects of OSN sensitization (due to receptor tuning) and reduced PN decay magnitude (putatively due to OSN population increases; see Discussion) on odor-evoked activity, which might synergize to promote sensitive and persistent long-range odor tracking toward the noni host fruit by *D. sechellia*, but not *D. melanogaster*.

We next pharmacologically impaired cholinergic neurotransmission to diminish excitatory connections of OSNs, which include OSN-PN and likely also OSN-LN synapses (Huang et al., 2010; Kazama and Wilson, 2008; Schlegel et al., 2021; Wilson, 2013). As expected, strong blockage essentially abolished odor-evoked PN responses (data not shown). More informatively, weak blockage led to enhanced decay in the VM5d PN responses of *D. sechellia* (**Fig. 6B**), as seen in untreated *D. melanogaster* (**Fig. 6B**). These observations suggest that excitatory neurotransmission from OSNs to PN and/or LNs also contributes to the temporal dynamics of PN responses. Consistent with this possibility, halving the number of OSN inputs in *D. sechellia* through removal of one antenna (OSN axons project to antennal lobes in both brain hemispheres (Schlegel et al., 2021)) enhanced the decay of this species’ VM5d PN responses (**Fig. 6C**). These data support the hypothesis that OSN number increase in *D. sechellia* modulates PN response dynamics.

## Discussion

Amongst the many ways in which animal brains have diverged during evolution, species-specific increases in the number of a particular neuron type are one of the most common. The genetic basis and physiological and behavioral consequences of such apparently simple changes have, however, remained largely unexamined. Here we have exploited an ecologically and phylogenetically well-defined model clade of drosophilids to study this phenomenon. We provide evidence that expansion of host fruit-detecting OSN populations in *D. sechellia* is a complex trait, involving contributions of multiple loci. Surprisingly, a larger number of OSNs does not result in sensitization of partner PNs nor in increased reliability of their responses. Rather we observed more sustained responses of PNs upon repetitive or long-lasting odor stimulation. While OSN number alone can influence the strength and persistence of odor-tracking behavior, this species-specific cellular trait is likely to synergize with increases in peripheral sensory sensitivity conferred by changes in olfactory receptor tuning properties (Auer et al., 2020; Dekker et al., 2006; Prieto-Godino et al., 2017) to enable long-distance localization of the host fruit of this species (**Fig. 6D**).

One important open question is how OSN population increases affect circuit properties and behavior. For one experimentally-accessible glomerulus, VM5d, we observed that PNs (which are unchanged in number between species) have a larger surface area and form more synapses with OSNs but show lower dendritic input resistance in *D. sechellia*. This anatomical and physiological compensation results in the voltage responses of PNs being very similar between species despite the increased OSN input in *D. sechellia*. Such compensation might reflect in-built plasticity in glomerular microcircuitry. Indeed, *D. melanogaster* shows intra-species difference in OSN number across glomeruli (Grabe et al., 2016) and the number of OSNs correlates with synapse number (**Fig. S13A** (Schlegel et al., 2021)). Moreover, a previous study in *D. melanogaster* characterized the consequence of (random) differences in PN numbers in a glomerulus (DM6): in glomeruli with fewer PNs (i.e. greater sensory convergence per neuron), individual PNs had larger dendrites, formed more synapses with OSNs, and exhibited lower input resistance (Tobin et al., 2017), analogous to our observations in *D. sechellia* VM5d.

Species-specific physiological responses to prolonged or repetitive odor stimuli likely involves multiple neuron classes. In *D. melanogaster*, adaptation of PNs to long odor stimuli occurs through lateral inhibition by GABAergic LNs (Nagel et al., 2015; Wilson and Laurent, 2005), as we confirmed here. However, such inhibition appears to be weaker in *D. sechellia* in pathways with more OSNs. Given the conserved total number of antennal OSNs between species, more OSNs in one pathway could lead to stronger inhibitory neurotransmission from the correspondingly larger glomerulus to other unchanged or smaller glomeruli. Critically, this could result in net weaker lateral inhibition from these glomeruli onto the expanded glomerulus, as seen in *D. sechellia*. We cannot exclude that LNs display species-specific innervations or connectivity, but testing this idea will require genetic drivers to visualize and manipulate subsets of this highly diverse neuron type (Chou et al., 2010). Finally, we note that it is also possible that differences exist in the intrinsic physiology of PNs due to, for example, differential expression of ion channels between species. Such differences – potentially unrelated to changes in OSN number – might be revealed by mining comparative transcriptomic datasets for these drosophilids (Lee and Benton, 2023).

Regardless of the precise mechanism, could more sustained PN responses convert to behavioral persistence? The main post-synaptic partners of PNs are lateral horn neurons and Kenyon cells, of which the latter (at least) have a high input threshold (Turner et al., 2008) for sparse coding. Assuming the threshold of these neurons is commensurate with maximum PN firing rate (which is higher in *D. melanogaster*), the relaxed decay in PN activity in *D. sechellia* might elongate downstream responses to persistent odors, which could drive valence-specific behaviors (Aso et al., 2014). Sensory habituation is advantageous for the brain to avoid information overload by attenuating constant or repetitive inputs and is a general feature across sensory modalities (O’Mahony, 1986). However, this phenomenon might be disadvantageous when navigating through sensory cues for a long period of time, for example, during olfactory plume-tracking. The reduced adaptation selectively in PNs in the expanded sensory pathways in *D. sechellia* that detect pertinent host odors provides an elegant resolution to this conflict in sensory processing, leaving conserved molecular effectors mediating synaptic neurotransmission and adaptation in other olfactory pathways intact.

Beyond *D. sechellia*, selective increases in ab3 neuron numbers have been reported in at least two other drosophilid species, which are likely to represent independent evolutionary events (Keesey et al., 2022; Linz et al., 2013) (**Fig. S13B**). Furthermore, Or22a neurons display diversity in their odor specificity across drosophilids (de Bruyne et al., 2010; Keesey et al., 2022). These observations suggest that this sensory pathway is an evolutionary “hotspot” where changes in receptor tuning and OSN population size collectively impact sensory processing, perhaps reflecting the potent behavioral influence of this pathway, as we have found in *D. sechellia*. More generally, given our demonstration of the important effect of OSN population size on host odor processing in *D. sechellia*, examination of how cell number increases modify circuit properties in other sensory systems, brain regions and species seems warranted.

## Acknowledgements

We thank Hokto Kazama, Johannes Larsch, Silke Sachse and members of the Benton laboratory for discussions and/or comments on the manuscript. We are grateful to Yoshi Aso, John Carlson, Sophie Caron, Tom Clandinin, Heather Dionne, Lukas Neukomm, Gerald Rubin, Stephan Sigrist, the Bloomington *Drosophila* Stock Center (NIH P40OD018537) and the Developmental Studies Hybridoma Bank (NICHD of the NIH, University of Iowa) for reagents. We also thank Jesse Isaacman-Beck for construct design, Anabela Rebelo Pimentel for her assistance with embryo alignments for microinjections, and the Lausanne Genomic Technologies Facility for sequencing. S.T. was supported by a Marie Skłodowska-Curie Actions Individual Fellowship (836783), an EMBO Long-Term Fellowship (ALTF 454-2019) and a Japanese Society for the Promotion of Science (JSPS) Overseas Research Fellowship (202360258). Research in J.R.A.’s lab was supported by the Swiss National Science Foundation (PP00P3_176956 and 310030_201188). C.F.R.W. was support by a National Institutes of Health grant (R01EY022638) to Tom Clandinin. F.v.B.’s laboratory was supported by the Air Force Office of Scientific Research (FA9550-21-0122). G.S. and J.M.J. were supported by NIH grants R01 DC018570 and R01 NS116584, the Richard and Susan Smith Family Award for Excellence in Biomedical Research, the Klingenstein-Simons Fellowship Award in Neuroscience, and an innovative research award from the Kavli Institute for Neuroscience at Yale University to J.M.J. T.O.A. and his laboratory were supported by a Human Frontier Science Program Long-Term Fellowship (LT000461/2015-L), a Swiss National Science Foundation Ambizione Grant (PZ00P3 185743) and the Fondation Pierre Mercier pour la science. R.B.’s laboratory was supported by the University of Lausanne, the Swiss National Science Foundation (310030B-185377) and ERC Consolidator (615094) and Advanced (833548) Grants.

## Author Contributions

S.T., T.O.A. and R.B. conceived the project. All authors contributed to experimental design, analysis and interpretation of results. Specific experimental contributions were as follows: S.T. performed all calcium imaging, photoactivation, tethered fly behavioral experiments and cloning of the *VT033006-LexA* construct. T.O.A. performed molecular biology experiments, generated transgenic lines, performed histology, Or85b QTL analysis, cell number quantifications and electrophysiology. G.S. performed PN electrophysiology, dye-filling and anatomical analysis, with guidance from J.M.J. L.A. performed molecular biology experiments, histology and quantifications of post-synaptic puncta. S.D.S. performed the wind tunnel behavioral experiments, with guidance from F.v.B. J.R.A. performed the Ir75b QTL analysis and contributed to the Or85b QTL analysis. L.L.P.-G. performed the Ir75b QTL analysis. D.L.S. contributed to the QTL analyses and generated the *UAS-CsChrimson* construct. S.C. contributed to the Ir75b QTL analysis. R.A.O. performed histology in the *D. melanogaster* subgroup. C.F.R.W. generated the parent *SPARC2-TNT* construct. S.T., T.O.A. and R.B. wrote the paper with input from all other authors. All authors approved the final version of the manuscript.

## Methods

### Data reporting

Preliminary experiments were used to assess variance and determine adequate sample sizes in advance of acquisition of the reported data. For electrophysiological recordings, data were collected from multiple flies on several days in randomized order. Within datasets, the same odor dilutions were used for acquisition of the data. The experimenter was blinded to the genotype for quantification of OSN numbers, and SPARC2-CsChrimson and SPARC2-TNT tethered fly assays, but not for other behavioral or physiological experiments.

### Drosophila strains

*Drosophila* stocks were maintained on standard wheat flour/yeast/fruit juice medium or, for those used in PN electrophysiology experiments, semi-defined culture medium (Backhaus et al., 1984) under a 12□h light:12□h dark cycle at 25°C. For all *D. sechellia* strains, a few g of Formula 4-24® Instant *Drosophila* Medium, Blue (Carolina Biological Supply Company) soaked in noni juice (Raab Vitalfood or Tahiti Trader) were added on top of the standard food. Wild-type, mutant and transgenic *Drosophila* lines used in this study are listed in **Supplemental Table 2.**

To generate *lozenge* (*lz*) *trans*-heterozygous *D. simulans/D. sechellia* hybrid flies, we group-aged 25-35 virgin males of *D. simulans* (*Dsim03 Or85b^GFP^*or *Dsim03 lz^RFP^;Or85b^GFP^*) for 6-7 days before combining them with 20-30 virgin females of *D. sechellia* (*Dsec07 lz^RFP^;Or85b^GFP^* or *Dsec07 Or85b^GFP^*). We lowered the fly food cap to restrict space and force interactions between the animals (which otherwise had a very low tendency to mate). Tubes were maintained at 22°C with strong light exposure and flipped every 3-4 days into a new tube. Progeny were collected and phenotyped 7-10 days post-eclosion.

### Constructs for CRISPR/Cas9-mediated genome engineering and transgenesis

*D. simulans Or85b*: for expression of a single sgRNA targeting the *D. simulans Or85b* locus, an oligonucleotide pair (**Supplemental Table 3**) was annealed and cloned into *BbsI*-digested *pCFD3-dU6-3gRNA* (Addgene #49410) as described (Port et al., 2014). To generate a donor vector for homologous recombination, homology arms (1-1.6 kb) were amplified from *D. simulans* (*Drosophila* Species Stock Center [DSSC] 14021-0251.195) genomic DNA and inserted into *pHD-Stinger-attP* (Auer et al., 2020) via restriction cloning. Both constructs were co-injected with a source of Cas9 (as described below) into *D. simulans* DSSC 14021-0251.003 and DSSC 14021-0251.004.

*D. simulans* and *D. sechellia lz*: to express multiple sgRNAs targeting the *lz* loci from the same vector backbone, oligonucleotide pairs (**Supplemental Table 4**) were used for PCR and inserted into *pCFD5* (Addgene #73914) via Gibson Assembly, as described (Port and Bullock, 2016). To generate donor vector for homologous recombination, homology arms (1-1.6 kb) were amplified from *D. sechellia* (DSSC 14021-0248.07) or *D. simulans* (DSSC 14021-0251.195) genomic DNA and inserted into *pHD-DsRed-attP* (Gratz et al., 2014) via Gibson Assembly. Species-specific constructs were co-injected with a source of Cas9 (as described below) into *D. simulans* DSSC 14021-0251.004 or *D. sechellia nos-Cas9* (Auer et al., 2020).

*D. sechellia UAS-SPARC2-D-CsChrimson*: we digested a *SPARC2-backbone* vector (Addgene #133562) with *SalI* and inserted a *CsChrimson-Venus* cassette after PCR amplification from *pBac(UAS-ChR2 CsChrimson,3xP3::dsRed)* via Gibson Assembly. The resulting *SPARC2-D-CsChrimson* cassette was amplified via PCR and inserted via restriction cloning into *pHD-3xP3-DsRed DattP-D. sechellia attP40* (this targeting vector for homologous recombination at the *D. sechellia attP40* equivalent site will be described in more detail elsewhere).

*D. sechellia UAS-SPARC2-D-TNT-HA-GeCO*: we first generated a *pHD-3xP3-DsRed_DattP-UAS-TNT-GeCO* vector by amplifying a *TNT-GeCO* cassette from *pHD-37.1_AttP5_LexAop_90_10_TNT-HA_p2A_jRGeCO1a* together with a *UAS* cassette and insertion into *pHD-3xP3-DsRed DattP* via Gibson Assembly. Subsequently, we transferred the *UAS-TNT-HA-GeCO* cassette into *pHD-3xP3-DsRed DattP-D. sechellia attP40* before assembling *pHD-3xP3-DsRed DattP-D. sechellia attP40 SPARC2-D-TNT-HA-GeCO* via Gibson Assembly (the GeCO calcium indicator was not used in the current study). Both *SPARC-D* transgenic lines in *D. sechellia* were generated via CRISPR/Cas9-mediated homologous recombination at the *attP40*-equivalent site in *D. sechellia*. To test the functionality of the *UAS-TNT-HA-GeCO* cassette, we also generated *UAS-TNT-HA-GeCO* transgenic lines via homologous recombination at the *attP40* locus in *D. melanogaster* and *D. sechellia*. However, in both species successful transformants did not survive pupariation, which was potentially due to low-level, Gal4-independent expression of the TNT effector.

*D. melanogaster VT033006-LexA*: to generate a *VT033006-LexA* construct, *pLexA-SV40-attB* was digested with *NotI*. A *VT033006* enhancer fragment was PCR-amplified from *pVT033006-Gal4-attB* (Tirian and Dickson, 2017). The insert and the linearized vector were joined by Gibson assembly. The vector was integrated into *D. melanogaster attP2* by BestGene Inc.

*D. sechellia VT033006-Gal4*, *VT033008-Gal4* and *VM5d-Gal4*: constructs carrying the *D. melanogaster* enhancer sequences (Tirian and Dickson, 2017) were integrated into *Dsec-white* (attP landing site on the X chromosome (Auer et al., 2020)) or *Dsec-attP40* (see next section). The *Dsec-nSyb-*Φ*C31* line was generated by integration of *nSyb-*Φ*C31* (Addgene *#*133868) into *Dsec-attP26* (see next section).

*D. sechellia UAS-myrGFP: pUAS-myrGFP, QUAS-mtdTomato(3xHA)* (Talay et al., 2017) was integrated into *Dsec-attP40*. *D. sechellia UAS-D*α*7:*GFP: flies were generated by P-element-mediated transgenesis of *p(UAS-D*α*7:GFP)* (Leiss et al., 2009) into *D. sechellia* DSSC 14021-0248.30 by WellGenetics.

*D. sechellia pBac(UAS-CsChrimson-Venus)*: we first amplified a *UAS-CsChrimson-Venus* cassette from *pUAS-ChR2 CsChrimson* (Klapoetke et al., 2014) and a *pBac* backbone (Horn and Wimmer, 2000) and combined both via Gibson assembly resulting in *pBac(UAS-CsChrimson-Venus)*. Subsequently, we digested *pBac(UAS-CsChrimson-Venus)* with *AscI*, amplified a *3xP3-DsRed* cassette (derived from gene synthesis, Genetivision) via PCR and combined both via Gibson Assembly resulting in *pBac(UAS-CsChrimson-Venus,3xP3-DsRed)*. Primer sequences for intermediate cloning steps are listed in **Supplemental Table 3**. PiggyBac-mediated transgenesis of *pBac(UAS-CsChrimson-Venus,3xP3-DsRed)* into *D. sechellia* DSSC 14021-0248.07 was performed in-house (see below) and the insertion site mapped to the third chromosome using TagMap (Stern, 2017). Beyond its use for optogenetic experiments, we took advantage of the visible 3xP3-DsRed marker to use the same line for introgression mapping (**Figure S2C,D**).

All plasmids were verified via Sanger sequencing before injection. Full details and oligonucleotide sequences are available from the corresponding authors upon request.

### *Drosophila* transgenesis

Except for specific constructs described above, mutagenesis/transgenesis of *D. sechellia*, *D. simulans* and *D. melanogaster* was performed in-house following standard protocols (Auer et al., 2020). For *piggyBac* and P-element transgenesis, we co-injected a *piggyBac* or *P-element* vector (300 ng µl^-1^) and *piggyBac* (Arnoult et al., 2013) or *P-element* helper plasmid (Stern et al., 2017) (300 ng µl^-1^). For CRISPR/Cas9-mediated homologous recombination, we injected a mix of an sgRNA-encoding construct (150 ng µl^-1^) and donor vector (500 ng µl^-1^) into *D. sechellia nos-Cas9* (Auer et al., 2020) or co-injected with *pHsp70-Cas9* (400 ng μl^−1^) (Addgene #45945; for *D. simulans* transgenesis) (Gratz et al., 2013). Site-directed integration into *attP* sites was achieved by co-injection of an *attB*-containing vector (400 ng µl^-1^) and *pBS130* (encoding ΦC31 integrase under control of a heat shock promoter (Addgene #26290) (Gohl et al., 2011)). The *Dsec-attP26* site (on chromosome 4) was generated via *piggyBac*-mediated random integration and *Dsec-attP40* via CRISPR-mediated homologous recombination and both will be described in more detail elsewhere. All concentrations are given as final values in the injection mix.

### Histology

Fluorescent RNA *in situ* hybridization (using digoxigenin- or fluorescein-labelled RNA probes) and immunofluorescence on whole-mount antennae were performed essentially as described (Saina and Benton, 2013; Silbering et al., 2011). Probes were generated using *D. sechellia* genomic DNA (*Or47a*, *Or88a*) and primers listed in **Supplemental Table 3**. Other published probes were either targeting *D. sechellia* (*Or42b*, *Or22a*, *Or85b*, *Or13a*, *Or98a*, *Or35a* (Auer et al., 2020)), *D. simulans* (*Or67a* (Auer et al., 2022)) or *D. melanogaster* transcripts (*Or56a*, *Or59b, Or9a*, *Or69aA* (Vosshall et al., 2000); *Or19a* (Couto et al., 2005); *Or83c*, *Or67d* (Chai et al., 2019)); all probes were used at a concentration of 1:50. Immunofluorescence on adult brains – with the exception of the visualization of dye-filled PN (described below) – was performed as described (Sanchez-Alcaniz et al., 2017).

The following antibodies were used: guinea pig α-Ir75b 1:500 (RRID:AB_2631093 (Prieto-Godino et al., 2017)), mouse monoclonal antibody nc82 1:10 (Developmental Studies Hybridoma Bank), rabbit α-GFP 1:500 (Invitrogen), and rat α-HA 1:500 (Roche). Alexa488-, Cy3- and Cy5-conjugated goat α-guinea pig, goat α-mouse, goat α-rabbit and goat α-rat IgG secondary antibodies (Molecular Probes; Jackson Immunoresearch) were used at 1:500.

### Image acquisition and processing

Except for dye-filled PN imaging and analysis (described below), confocal images of antennae and brains were acquired on an inverted confocal microscope (Zeiss LSM 710) with an oil immersion 40× objective (Plan Neofluar 40× Oil immersion DIC objective; 1.3 NA). For quantification of synapse numbers, images were taken using a 63× objective (Plan-Apochromat 63× Oil immersion DIC M27; 1.4 NA) with a zoom of 3×, centering the image on the glomerulus of interest. Images were processed in Fiji (Schindelin et al., 2012). *D. sechellia* brains were imaged and registered to a *D. sechellia* reference brain (Auer et al., 2020) using the Fiji CMTK plugin (https://github.com/jefferis/fiji-cmtk-gui).

Cell number quantification: the number of OSNs expressing a specific *Or* was quantified using Imaris (Bitplane) or the Fiji Cell Counter tool. For GFP-expressing Or85b OSNs for the QTL analysis, we imaged GFP and, in the 568 nm channel, cuticular autofluorescence. After subtraction of the cuticular fluorescence signal from the GFP signal using the Subtraction tool in Fiji, we quantified the number of GFP-positive nuclei using the Surface Detection tool in Imaris. For Ir75b neurons, we found that α-Ir75b immunofluorescence resulted in labelled cells having a range of intensities. To ensure that the cell quantifications were reproducible, counting was performed manually by three experimenters. Images resulting in disagreements were re-checked and either resolved or removed from the analyses.

Glomerular and synapse quantification: glomerular volumes were calculated following segmentation with the Segmentation Editor plugin of Fiji using the 3D Manager plugin. The number of post-synaptic sites per glomerulus was quantified in Imaris as described (Mosca and Luo, 2014), setting punctum size to 0.45 μm^3^ for all images.

### Quantitative trait locus mapping

*Or85b phenotyping:* flies expressing nuclear-localized GFP in Or85b neurons were placed individually into 96-well plates whose bottom was replaced by a metal mesh. Antennae were removed via shock-freezing in liquid nitrogen and collected in 4% paraformaldehyde-3% Triton-PBS as described (Saina and Benton, 2013). After 3 h of fixation, antennae were washed twice in 3% Triton-PBS and twice in 0.1% Triton-PBS and mounted in Vectashield (Vectorlabs) on 30-well PTFE printed slides (Electron Microscopy Sciences) before imaging on a Zeiss LSM710 confocal microscope. The fly bodies were transferred (by inversion) into separate 96-well plates and frozen at -20°C. Genomic DNA of individual flies was extracted using the ZR-96 Quick-gDNA MiniPrep kit (Zymo Research).

*Ir75b phenotyping*: flies were processed as described above. For antennae, after washing in 0.1% Triton-PBS, Ir75b immunofluorescence was performed. Antennae were mounted in Vectashield (Vectorlabs) on 30-well PTFE printed slides (Electron Microscopy Sciences) before imaging on a Zeiss LSM710 confocal microscope.

*Sequencing and genotyping*: genomic DNA of individual flies was tagmented with in-house produced Tn5 as described (Picelli et al., 2014). In brief, Tn5 was charged with adaptors and mixed at a concentration of 5 ng μl^-1^ with 1 μl of genomic DNA. After tagmentation, Tn5 was de-activated by addition of 0.2% sodium dodecyl-sulphate and sample specific sequencing adaptors were added by PCR amplification. The resulting PCR amplicons were cleaned-up with AMPure XP bead-based reagents (Beckman Coulter Life Science), DNA concentration and fragment distribution quantified on a fragment analyzer (Agilent) and single-end sequenced on an Illumina HiSeq sequencer.

*Data analysis*: to align sequencing reads, the parental genomes dsim r2.02 and dsec r1.3 were used as reference. Introgressed genomic regions were inferred using MSG software (http://www.github.com/JaneliaSciComp/msg). Output of MSG was thinned using the “pull_thin” utility (http://www.github.com/dstern/pull_thin) and read into R (v4.4.1) using the “read_cross_msg” utility (http://www.github.com/dstern/read_cross_msg). QTL mapping was carried out with the qtl package (v1.5) using the Haley-Knott method and significance was determined using 1000 permutation tests (n.perm = 1000). For the Or85b mapping experiment, tests for interactions between the QTL on chromosomes 3 and X were performed using the “fitqtl” function.

### Two-photon calcium imaging

*Animal preparation*: flies of the appropriate genotype (described in the respective figure legends) were collected (females and males co-housed) and reared in standard culture medium (see above, with an addition of blue food with noni juice for *D. sechellia*). Female flies aged 5-8 days after eclosion were used for the experiments.

*Sample preparation*: flies were anaesthetized by placing them into an empty vial that was cooled on ice for no longer than ∼10 min. Further steps were performed under a dissection microscope, adapting a previous protocol (Silbering et al., 2012). A small drop of blue light-curing glue (595987WW, Ivoclar Vivadent) was placed on top of the copper grid (G220-5, Agar Scientific). Single flies were introduced into the mounting block fixing the back of the fly head to the copper grid with the curing glue. The fly head was slightly bowed down with forceps (to achieve antennal lobe imaging from the dorsal side) while the blue curing light (bluephase C8, Ivovlar Vivadent; placed at least 1 cm from the fly to avoid tissue damage) was focused. We avoided using the wire that was previously used to pull down the antennal plate (Auer et al., 2020; Silbering et al., 2012), as we found that it very frequently damaged the antennal nerves and therefore disrupted central olfactory responses, particularly in *D. sechellia*; the cactus spine and screw previously used to immobilize the fly head were also no longer necessary. An antennal shield (Silbering et al., 2012) was placed over the top of the fly head with the hole positioned centrally, fixed with beeswax to the top of the mounting block. Two-component silicon (Kwik-Sil, World Precision Instruments) was mixed with a toothpick and poured into the hole of the antennal shield to seal the gap between the plate and the fly head, avoiding any leakage onto the antennae. As the silicon was left to harden (during 10-15 min), we used blunt forceps to gently remove silicone on the cuticle on top of the head capsule. A drop of adult hemolymph (AHL) saline (108 mM NaCl, 5 mM KCl, 2 mM CaCl_2_, 8.2 mM MgCl_2_, 4 mM NaHCO_3_, 1 mM NaH_2_PO_4_, 5 mM trehalose, 10 mM sucrose, 5 mM HEPES, pH 7.5) was added into the hole of the antennal shield. Using a blade-splitter, a small rectangular hole was cut in the head cuticle between the eyes and above the antennal plate. Tracheae and glands above the brain were removed with fine forceps. Finally, the brain was rinsed with AHL saline at least 3 times until the antennal lobe appeared clear under the dissection microscope.

*Odorant preparation*: serial dilutions of odors were prepared in a fume hood. The solvents used for odor dilutions were different between glomeruli, as some glomeruli were extremely responsive to particular solvents: Or85b/VM5d (*2*- heptanone (CAS 110-43-0) in dichloromethane (DCM)), Or22a/DM2 (methyl hexanoate (CAS 106-70-7) in paraffin oil), Or59b/DM4 (methyl butyrate (CAS 623-42-7) in DMSO (for dose-responses) or paraffin oil (for pulsed stimuli)), Or92a/VA2 (*2*,*3*-butanedione (CAS 431-03-8) in paraffin oil). Or85b/VM5d neurons were especially sensitive to odorant contamination, so we prepared a new set of *2*-heptanone dilutions when solvent responses started to become evident (approximately every two weeks). Furthermore, before preparing *2*-heptanone and methyl butyrate, we washed the odor-containing vial, lid and pipetting tips with DCM and DMSO, respectively. As DCM is highly volatile at room temperature, *2*-heptanone/DCM odor dilutions were stored at 4°C.

*Image acquisition*: images were acquired using a commercial upright two-photon microscope (Zeiss LSM 710 NLO). An upright Zeiss AxioExaminer Z1 was fitted with a Ti:Sapphire Chameleon Ultra II infrared laser (Coherent) as excitation source. Images were acquired with a 20× water dipping objective (Plan-Apochromat 20× W; NA 1.0), with a resolution of 128×128 pixels (0.8926 pixels µm^-1^) and a scan speed of 6.30 µs pixel^-2^ (for one-time odor stimulation) or 3.15 s pixel^-2^ (for pulse train odor stimulation). The excitation wavelength was set to 930 nm. The output power was modified according to the baseline fluorescence of GCaMP6f, which varied substantially between animals (except for pharmacology experiments, where laser output was consistent to enable comparison of raw fluorescence across animals). The power was set such that the baseline fluorescence was above the detection limit, and that the maximum fluorescence was below saturation, and thereafter unchanged for a given animal. Emitted light was filtered with a 500-550 nm band-pass filter, and photons were collected by an internal detector. Each measurement consisted of 50 images acquired at 4.17 Hz (for one-time odor stimulation) or 8.34 Hz (for pulse train odor stimulation), with stimulation starting ∼5 s after the beginning of the acquisition and lasting for 1 s (for one-time odor stimulation) or 200 ms followed by a 200 ms interval repeated ten times (for pulse train odor stimulation).

*Olfactory stimulation*: antennae were stimulated using a custom-made olfactometer (Auer et al., 2020; Silbering et al., 2012). In brief, antennae were permanently exposed to air flowing at a rate of 1.5 l min^-1^ by combining a main airstream of humidified room air (0.5 l min^-1^) and a secondary stream (1 l min^-1^) of normal room air. Both air streams were generated by vacuum pumps (KNF Neuberger AG) and the flow rate was controlled by two independent rotameters (Analyt). The secondary airstream was guided either through an empty 2 ml syringe or through a 2 ml syringe containing 20 µl of odor or solvent on a small cellulose pad (Kettenbach GmbH) to generate odor pulses. To switch between control air and odor stimulus application, a three-way magnetic valve (The Lee Company, Westbrook, CT) was controlled using MATLAB via a VC6 valve controller unit (Harvard Apparatus). The order of the odor stimuli was always from lower to higher concentrations, preceded by the solvent control. Successive odor stimulations were separated by 1 min intervals.

*Pharmacology*: for pharmacological experiments, drugs were diluted in AHL saline to the following final concentrations: 100 µM Picrotoxin (P1675-1G, Sigma-Aldrich, CAS 124-87-8), 50 µM CGP54626 hydrochloride (1088/10, TOCRIS, CAS 149184-21-4), 200 µM (low dose) or 2 mM (high dose), mecamylamine hydrochloride (M9020-5MG, Sigma-Aldrich, CAS 826-39-1). Drugs were applied to the fly after normal recording (“saline / naïve”) by exchanging the AHL saline with drug-diluted saline five times and incubating the preparation for a further 15-40 min before performing further recordings. For mecamylamine application experiments, samples were subsequently washed with AHL saline five times and incubated for 15-40 min, followed by another recording session (“Washed-out”). The lens was meticulously washed with ultrapure water between each session.

*Data analyses*: data were processed using Fiji and custom written scripts in R. First, the image stacks were passed through the StackReg plugin (Thevenaz et al., 1998) (transformation: Rigid Body) to correct for movement artefacts. Using Fiji, a circular region of interest (ROI) was set within the glomerulus of interest on the left half of the brain image (except when the signal was weak, in which case the right half of the image was used). The signal intensity averaged across the ROI for each timeframe (hereafter *F*) was used to calculate the normalized signal 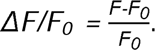 Here, *F_0_* (baseline fluorescence) was calculated as the average *F* during frames 16-19 (1 s before olfactory stimulus onset). The peak Δ*F/F_0_* value (which represents the odor response intensity) was calculated as the maximum Δ*F/F_0_* value during frames 20-23 (1 s during olfactory stimulation). We noticed that the maximum Δ*F/F_0_* value itself was often very different between species/genotypes, presumably due to different expression levels of GCaMP6f. This should, in theory, not affect the normalized Δ*F/F_0_* value, but we did observe a saturation of neuronal responses above a certain odor concentration, even if the peak Δ*F/F_0_* value was lower than in other species/genotypes. To compare the dose-response effect between species and genotypes, we further introduced a normalization step. Normalized peak response for each odor dilution 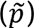 was calculated as follows: 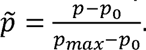 Here, *p* denotes peak Δ*F/F_0_* value of a given dilution (median value across animals), *p_max_* denotes maximum *p* among all the dilutions, and *p_o_* denotes *p* from the minimum response. Thus, the normalized peak response 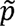 takes a value between 0 and 1, where 0 means absence of odor responses and 1 means saturation. This step allowed us to compare the dose-response curve based on the relative response within species and genotypes, regardless of the absolute Δ*F/F_0_* value. Dynamic ranges were quantified as *p_max_ - p_o_* in each animal rather than taking the median values across animals.

### Photoactivation

*Animal preparation*: flies of the appropriate genotype (see figure legends) were collected, reared and prepared in the same way as for the two-photon calcium imaging. Female flies aged 6-9 days after eclosion were used for the experiments.

*Photoconversion and image acquisition*: the hardware setup was the same as in two-photon calcium imaging. Glomerular location was identified by a brief 930 nm scan of the entire antennal lobe. An oval ROI was placed inside the glomerulus of interest, where the ROI was made small enough so that the movement during photoconversion did not result in non-specific labelling. Where non-specific labelling was observed after imaging, data were excluded from further analyses. During photoconversion, the resolution was set to 512×512 pixels (3.5704 pixels µm^-1^) and the scan speed to 0.79 µs pixel^-2^. Excitation wavelength was shifted to 760-780 nm with a power output of 5-15%. Photoconversion was performed by scanning inside the ROI repeatedly (in a single z-plane) for ∼15 min. Around 5 min after the beginning of the session, a brief 930 nm scan was performed to check the conversion efficacy and specificity. If photoconversion was weak at this point, we increased the power output. The sample was then placed in a humidified chamber for 30-60 min to allow the diffusion of photoconverted C3PA-GFP. Finally, the fly’s body was removed to reduce motion artefacts and the sample placed again under the two-photon microscope. For imaging, the resolution was set to 1024×1024 pixels (2.3803 pixels µm^-1^) and the scan speed to 1.58-3.15 µs pixel^-2^. The excitation wavelength was shifted back to 930 nm with the power output adjusted to enable visualization of neurite processes. Z-stack images were obtained with a spacing of 1 µm.

### Electrophysiology

*Single sensillum recordings*: single sensillum electrophysiological recordings were performed essentially as previously described (Auer et al., 2020), using 5-7 day-old female flies, which were grown on standard medium mixed with 0.2 mM all-*trans* retinal (Klapoetke et al., 2014), and, for *D. sechellia*, addition of 10% noni juice. Optogenetic stimulation was performed by exposing one antenna with increasing light intensities via an optic fiber as described in the **Tethered fly assay** section (see below). In SPARC2 experiments, ab3 sensilla were identified by location and the use of diagnostic odors. For the data shown in **Fig. 3C**, light-sensitive (experimental group) and non-responding neurons (control) were analyzed. Corrected responses were calculated as the number of spikes in a 0.5 s window from the beginning of illumination, subtracting the number of spontaneous spikes in a 0.5 s window 2 s prior to illumination, and multiplying by 2 to obtain spikes s^−1^. Recordings were performed on a maximum of three sensilla per fly. Exact *n* values and mean spike counts for all experiments are provided in **Supplemental Table 1**.

*Whole-cell patch clamp recordings:* for *in vivo* VM5d PN recordings, flies were prepared and dissected as previously described (Jeanne and Wilson, 2015). Female flies aged 1-2 days post-eclosion were used for the experiments; one neuron was recorded per brain. The internal patch pipette solution contained 140 mM potassium aspartate, 10 mM *4*-(*2*-hydroxyethyl)-*1*-piperazineethanesulfonic acid, 4 mM MgATP, 0.5 mM Na_3_GTP, 1 mM ethylene glycol tetraacetic acid, 1 mM KCl and, for cell labelling described below, 13 mM biocytin hydrazide. The pH was adjusted to 7.3, and the osmolarity was adjusted to ∼265 mOsm. The external saline contained 103 mM NaCl, 3 mM KCl, 5 mM *N*-tris(hydroxymethyl) methyl-*2*-aminoethane-sulfonic acid, 8 mM trehalose, 10 mM glucose, 26 mM NaHCO_3_, 1 mM NaH_2_PO_4_, 1.5 mM CaCl_2_, and 4 mM MgCl_2_. The osmolarity was adjusted to 270-273 mOsm, and the saline was bubbled with 95% O_2_ and 5% CO_2_ and reached an equilibrium pH of 7.3. Saline was continuously superfused over the fly during recording. Recordings were acquired with an Axopatch 700B or 200B model amplifier, low-pass filtered at 4 or 5 kHz, and digitized at 10 kHz. Patch pipettes, made from borosilicate glass, were pressure-polished. The estimated final pipette tip opening was less than 1 μm in diameter, and the pipette resistance was 15-45 MΩ.

*Olfactory stimulation:* serial dilutions of *2*-heptanone (in mineral oil) were freshly prepared before each experiment. A custom-made olfactometer was used to deliver odor to flies. Antennae were consistently exposed to a stream of air at 363 ml min^-1^. Another stream of air (5.3 ml min^-1^) was directed through a solenoid valve into a 2 ml vial (Thermo Scientific, National C4011-5W) containing either mineral oil alone or an odor solution in mineral oil. Odor delivery, controlled by a custom MATLAB script and a three-way solenoid valve (The Lee Company, Westbrook, CT), lasted 2 s. The series of stimuli always started with the solvent control followed by increasing odor concentrations. Custom-written MATLAB scripts were used to detect spikes based on the first derivative of the voltage trace. A threshold was set for each recording and all spikes were visually inspected to eliminate both false positive and false negative detections. The spike time was defined as the time of the peak of the first derivative of the voltage waveform. An average of three trials was taken for each concentration for each cell. Corrected responses were calculated as the spike rate in a 50 ms (**Fig. 5E and Fig. S7B**) or 500 ms (**Fig. S7B**) window and subtracting the spontaneous spike rate (computed in a 500 ms window 2 s prior to stimulation). Exact *n* values and mean spike counts for all experiments are provided in **Supplemental Table 1**.

*Projection neuron backfilling and reconstruction*: each brain was dissected out of the head capsule after the recording and fixed with 4% PFA (w/v) in PBS for 20 minutes at room temperature. After washing with PBS-T [PBS with 0.2% (v/v) Triton X-100 (Sigma Aldrich, #X100)], brains were incubated with streptavidin Alexa Fluor 568 (1:1000) (Invitrogen S11226) and nc82 (1:50) and 10% Normal Goat Serum in 0.2% PBS-T overnight at RT. The brains were washed and incubated with streptavidin Alexa Fluor 568 (1:1000) and anti-mouse Alexa Fluor 633 (1:500) and 10% Normal Goat Serum in 0.2% PBS-T overnight. After the final wash with PBS, brains were mounted in Vectashield H-1000 (Vector Laboratory, Burlingame, CA) anti-fade mounting medium for confocal microscopy. Brains were imaged with a Zeiss LSM880 confocal microscope. Confocal images displayed Biocytin fills in all recordings, and dendritic surface area was measured using IMARIS 10.0.0 (Bitplane) through a semi-automated generation of surfaces for each dendritic arbor.

For one *D. melanogaster* recording, the biocytin fill revealed two coupled cells: one innervating the VM5d glomerulus and the other innervating a different glomerulus. Correspondingly, two distinct spike waveforms were clearly discernible in the voltage trace. We excluded this recording from membrane voltage and input resistance analyses but included it in spike rate analysis because the VM5d PN spikes could be identified by their clear responses to *2*-heptanone. The VM5d arbor in this fill was included in the dendritic morphology analysis.

### Tethered fly assay

*Assay*: the assay was built on a solid breadboard (Thorlabs). The fly tether was made by inserting and gluing an insect pin (Austerlitz, φ = 0.20 mm) to a 200 µl pipette tip, which was mounted on a magnetic articulated stand (NOGA). Two microphones (lavalier microphone, RODE) were placed ∼1 mm from the tip of the wings of the fly, connected to a USB audio interface (Rubix 22, Roland) via TRS-XLR adaptors (VXLR, RODE). The audio interface was connected to Raspberry Pi computer (Raspberry Pi 4 1.5 GHz Quad-Core, 8GB RAM), which ran the real-time feedback program (described below) based on the acoustic inputs. The output of the feedback system was SPI-connected to a DotStar LED strip (1528-2488-ND, Adafruit; cut to 30 LEDs), which was bent to make a U-shape that covers >180° of the fly’s horizontal view. The spatial frequency of the visual guide was set to ∼0.036 mm^-1^ (one illumination in every 4 LEDs). The PTFE odor port was placed at ∼30° from the right side, facing the fly ∼1 cm apart to provide unilateral olfactory stimulation. The suction port was placed at the opposite end of the odor port, ∼2 cm from the fly, to stabilize the odor plume. For optogenetic experiments, single antennae were illuminated using a custom-made optic fiber (G050UGA, Tubing: FT030, End 1: SMA, End 2: Flat Cleave, Thorlabs), of which the cleaved end was placed ∼0.1 mm from one antenna. The position of the optic fiber was adjusted using a 3D microcontroller (UM-3C, Narishige), and was monitored by a Pi NoIR camera prior to each session. The fiber was connected to 660 nm fiber-coupled LED (M660FP1, Thorlabs) via a compact LED driver (LEDD1B, Thorlabs).

*Real-time feedback system*: the feedback system was run by a custom Python script. The recording was performed binocularly from each wing at the rate of 44,100 Hz. Each session (20 s) was divided into 0.1 s intervals. The raw wing beat amplitude (*r_[L]_, r_[R]_*) was defined by the difference between the maximum and minimum sound amplitude within each interval. The raw wing beat was filtered by calculating the median across the three most recent intervals (*r_f[L]_, r_f[R]_*). Prior to the experimental session, each fly was calibrated by a “mock” session, where the visual guide was fixed and the flies beat their wings to obtain the mean (*μ_mock[L]_, μ_mock[R]_*) and SD (*σ_mock[L]_, σ_mock[R]_*) of the filtered wing beat amplitude as well as the SD of the difference between the left and right wing beat amplitudes (*σ_mock[L-R]_*). In the experimental sessions, the filtered raw wing beat amplitudes were standardized using the mean and SD obtained in the mock session 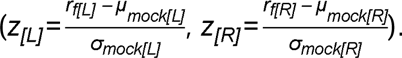 The Δ*WBA* was defined by *z_[L]_ -z_[R]_* as a readout of turning behavior. In each interval, the visual guide was rotated counter-clockwise (shifting by one adjacent LED) if *ΔWBA > 3σ_mock[L-R]_* and clockwise if clockwise if *ΔWBA < 3σ_mock[L-R]_*. The *z_[L]_, z_[R]_* values were saved after each session for downstream analyses.

*Animal preparation*: flies of the appropriate genotype were collected (females and males co-housed). For odor response experiments, females 5-9 days after eclosion were used (except for SPARC2-D-TNT experiments, where flies were 0-1 day old since the experimental group did not survive for long). For experiments with apple cider vinegar (Migros, M-Classic), flies were starved for 5-7 h. Fly rearing for optogenetic experiments is as described in **Electrophysiology** section. Flies were reared in retinal-containing food for 6-7 days, and females 6-8 days after eclosion were used.

*Sample preparation*: flies were anaesthetized on ice and attached to the fly tether using blue-curing glue. For optogenetic experiments, fly forelegs were cut to prevent the flies from perturbing the optic fiber. The fly tether was mounted on a magnetic stand, and the fly positioned centrally and equidistantly between the microphones. For optogenetic experiments, videos were taken during positioning of optic fibers to confirm that the illumination (at intensity 3, as described below) was confined to one antenna. For SPARC2-D-TNT-HA experiments, as transgene expression was not detected in a fraction of flies (for unknown reasons), post-hoc HA immunofluorescence was performed in individual antennae and/or brains and the behavioral data for animals with positive labelling in OSN cell bodies and/or axon termini were retained in the downstream analyses.

*Olfactory stimulation*: odorants (5 ml) were contained in 15 ml Falcon tubes with two syringe needles pierced at the lid. Tubes were connected to the syringe needle outlets to provide odor stimulation from the headspace. The flies were permanently exposed to water vapor at a rate of 0.5 l min^-1^. A three-way magnetic valve (The Lee Company, Westbrook, CT) was controlled using MATLAB via a VC6 valve controller unit (Harvard Apparatus) to switch the airflow from water to odorants. Either 10 consecutive pulses of 500 ms with 500 ms intervals or a single 10 s pulse were used as olfactory stimuli. Noni juice was diluted in water while benzaldehyde was diluted in paraffin oil.

*Optogenetic stimulation*: LED illumination was controlled by sending TTL signals to the LED driver. 10 consecutive pulses of 500 ms with 500 ms intervals were used. The intensity of the illumination was modified by the dial on the LED driver, which could be modulated from intensity 0 (mock-stimulation) to 6 (maximum). Intensity 4 was used for stimulation in most experiments, except for low-intensity stimulation (intensity 3) to match the OSN spike rate with SPARC2-D-CsChrimson experiments. For SPARC2-D-CsChrimson experiments, we noticed that the animals either displayed expression of CsChrimson (as detectable by the Venus tag) in about half of the Or22a OSN population or had no expression at all. The reason for this heterogeneity is unknown but we performed post-hoc imaging of Venus fluorescence in individual antennae of all animals to analyze data from only those that expressed the transgene.

*Data analyses*: data were analyzed with custom programs in R. The quantification was performed in 1 s time windows corresponding to pre-stimulus baseline (4-5 s) and individual odor-pulse responses (1^st^ peak: 5-6 s, 2^nd^ peak: 6-7 s, …, 10^th^ peak: 14-15 s). For quantification of Δ*WBA* in each time window, the Δ*WBA* values above the 50% quantile were used, to avoid picking outliers and non-attractive epochs.

### Wind tunnel assay

Free flight tracking was performed in a 1 m × 0.5 m × 0.5 m wind tunnel housed in a temperature- and humidity-controlled room (22°C, 60% RH). The wind tunnel floor was illuminated by blue LEDs underneath a light-diffusive film that lined the tunnel floor, overlaid with a grid of infrared transmissible film; this grid formed a checkerboard-like lattice on the floor to provide flies with ventral optic flow information. The walls of the wind tunnel tapered from blue on the floor to black along the ceiling. Along the top of the wind tunnel were two white LED strips, to provide sufficient orange light for *Drosophila* photoreceptor re-isomerization (Byk et al., 1993). Total illumination in the wind tunnel was 452 Lux as measured from the wind tunnel center (LX1330B light sensor, Dr. Meter).

Fly tracking was performed with 12 synchronized Basler acA720 cameras (Basler AG, Ahrensburg) recording at 100 Hz (**Fig. S4A**). The walls and floor of the wind tunnel were homogenously illuminated with arrays of infrared LEDs, and cameras were outfitted with IR pass filters to track flies using only infrared wavelengths. Tracking was performed using Braid software (Straw et al., 2011). Odor plumes were introduced in the wind tunnel through an odor port constructed from a 20 cm tall rigid acrylic tube that had a 90° bend towards the top so that its outward facing opening was parallel to the wind (**Fig. S4B**). The odor port was installed on the far upwind end of the tunnel and its open face was covered by an aluminum mesh to prevent flies from entering the odor plumbing. Plumes were generated by fluxing air through a mass flow controller (Alicat Scientific, Tuscon) at 200 cm^3^ min^-1^. The air was then bubbled into 150 ml noni juice at the bottom of a jar and subsequently through tubing leading to the odor port in the wind tunnel.

For wind tunnel experiments we used 3-7 day old female flies. 15 animals were collected in the morning and starved by placing them into a tube containing a moist Kimwipe for 8 h. The flies were placed in the wind tunnel with a 40 cm s^-1^ air flow and tracking initiated. Flies were allowed to fly about the wind tunnel volume for 16-20 h before the recording was terminated the following day.

*Data analyses*: data were analyzed using Python (v3.11). We first filtered out all trajectories from flies that were walking on the tunnel floor or ceiling to focus analyses on flying animals. Our trajectory inclusion criteria were: (i) >500 ms long; (ii) >10 cm in the horizontal plane; (iii) a median position of >5 cm away from any of the tunnel walls; (iv) passed within a 10 cm radius of where the odor plume was aligned in the y-z plane (this excluded flies that simply transited the wind tunnel along the ceiling, far away from the plume).

To analyze the radial distance from the plume, we binned the point cloud generated by all trajectories into 5 cm thick cross-sections, beginning 5 cm downwind of the plume source. We then pared the point cloud further to 10 cm above and below the plume (0.1<z<0.3) to isolate trajectory portions that were most likely interacting with the plume (**Fig. S4E**), for analyses with or without this restriction. Based on the occupancy maps of trajectories in space from both wild-type *D. sechellia* and *D. melanogaster*, we assumed that the plume sank by 5 cm from the odor nozzle to the end of the wind tunnel and calculated the plume centerline using this mode (for comparisons of straight versus sloped plume models see **Fig. S4E-F**). From each of the segmented point clouds we report the mean as the mean of all points, independent of trajectories that contributed to them. Results were robust to whether we instead analyzed each trajectory as an independent sample. For course direction distributions, we included only the point cloud within a 3 cm radius of our estimated plume volume and calculated the kernel densities of the course direction of all points within the estimated plume volume. We further divided the data to the point cloud in the downwind half of the wind tunnel, where trajectories typically began, and the upwind half of the wind tunnel, terminating 5 cm downwind of the plume’s origin.

## Statistics and reproducibility

Data were analyzed and plotted using Excel, R (v3.2.3; R Foundation for Statistical Computing, Vienna, Austria, 2005; R-project-org), MATLAB (2023a), GraphPad Prism (10.1.1) and Python (v3.11).

## Data, code and biological material availability

All relevant data supporting the findings of this study are included as source data or available from the corresponding authors upon request. Code used for analyses and all unique biological materials generated in this study are available from the corresponding authors upon request.

## Supplemental Figure Legends

**Figure S1: Investigation of potential mechanisms underlying ab3 sensilla expansion.**

**(A)** Quantification of *Or22a/(b)* RNA expressing OSNs in the antenna of six *D. sechellia*, *D. simulans* and *D. melanogaster* strains. For strain details see **Supplemental Table 2**. Wilcoxon signed-rank test, *P* values adjusted for multiple comparisons using the Benjamini and Hochberg method. Comparisons to *Dsec07* (genetic background of transgenic lines) are shown. NS, not significant (*P* > 0.05); \*\**P* < 0.01; \*\*\**P* < 0.001.

**(B)** Top, immunofluorescence for GFP and RNA FISH for *Or85b* on whole-mount antennae from *D. melanogaster Or49b-GFP* animals. Arrowheads indicate paired neurons. Or85b neurons are housed in ab3 (with Or22a neurons) and ab6 sensilla (with Or49b neurons), but only the ab3 population is expanded in *D. sechellia* (Fig. 1C). Scale bar, 25 µm.

**(C)** Comparison of the number of *Or22a* RNA-expressing OSNs in the antenna of *D. sechellia* raised on (left) and without (right) noni supplement. Scale bar, 25 µm. Quantification to the right. Wilcoxon rank-sum test. NS, *P* > 0.05.

**(D)** Top left, schematic of the ab2 sensillum housing Or59b and Or85a neurons. Top right, RNA FISH for *Or59b* on whole-mount antennae from *D. melanogaster, D. simulans and D. sechellia* wild-type animals (females). Scale bar, 25 µm.

**(E)** Top, schematic of developmental transitions from larval antennal disc to adult antenna. Schematic of the larval antennal disc, within which concentric arcs of sensory organ progenitors (SOPs) are specified, each of which gives rise to a sensillum. Different SOP types have a stereotyped developmental origin in a given arc (adapted from (Chai et al., 2019)). Arrows indicate the change in relative population size in *D. sechellia* compared to *D. melanogaster*. No obvious relationship between ab3 number increase and compensatory reduction in sensilla derived from SOPs in the same or neighboring arcs is evident.

**(F)** Representative pictures of the reciprocal hemizygosity test at the *Or22a/b* and *Or85c/b* loci using RNA FISH results shown in Fig. 1D. Schematics on top indicate expression from the respective alleles.

**Figure S2: Genetic analysis of OSN cell number expansion.**

**(A)** Left, schematics depicting the introduction of a nuclear-localized GFP reporter (GFPnls) at the *Or85b* locus of *D. sechellia* and *D. simulans* via CRISPR/Cas9 genome engineering. Right, GFP signal in antennae of *DsecOr85b^GFP^* and *DsimOr85b^GFP^* animals. Scale bar, 25 µm.

**(B)** Immunofluorescence for GFP and *Or22a* RNA FISH on whole-mount antennae from *DsecOr85b^GFP^* animals. Arrowheads indicate neighboring neurons. Scale bar, 25 µm.

**(C)** Introgression of chromosomal fragments marked with a transgenic RFP marker (dashed line) from *D. sechellia* into *D. simulans* spanning varying extents of the QTL peak on chromosome 3.

**(D)** Left, quantification of GFP-expressing neurons in F2 backcrosses of the *D. sechellia* RFP transgenic line to *D. simulans Or85b^GFP^* flies comparing RFP positive and negative siblings (females only). Right, quantification of GFP-expressing neurons in the five *D. sechellia* introgression lines depicted in (**A**) comparing RFP positive and negative siblings. *n* are listed in the figure. Wilcoxon signed-rank test. NS, not significant (*P* > 0.05); \**P* < 0.05; \*\*\**P* < 0.001.

**(E)** Left, location of the *lozenge* (*lz*) gene (dashed line) relative to the QTL peaks detected on the X chromosome. Right, schematics depicting the *lozenge* gene organization and the structure of mutant alleles in both *D. sechellia* and *D. simulans*. The fluorescent marker was integrated into the first coding exon.

**(F)** Comparison of antennal morphology in wild-type and *lz* mutant *D. sechellia* and *D. simulans*. In both species, loss of *lz* results in a lack of basiconic sensilla (white arrowheads) compared to wild-type, similar to *D. melanogaster lz* mutants (Gupta et al., 1998). Scale bar, 25 µm; inset scale bar, 5 µm.

**(G)** Quantification of GFP-expressing neurons in *Dsimlz^RFP^* heterozygote mutant and wild-type siblings (females only). Wilcoxon signed-rank test. \*\**P* < 0.01.

**(H)** Quantification of GFP-expressing neurons in *Dseclz^RFP^* heterozygote mutant and wild-type siblings (females only). Wilcoxon signed-rank test. \*\*\**P* < 0.001.

**(I)** Quantification of GFP-expressing neurons in *trans*-heterozygote hybrid siblings (females only) carrying either a *Dseclz^RFP^* or *Dsimlz^RFP^*allele. Wilcoxon signed-rank test. NS, not significant (*P* > 0.05).

**(J)** Quantification of Ir75b neurons in wild-type *D. sechellia* (DSSC 14021-0248.25) and *D. simulans* (DSSC 14021-0251.195) (mixed genders) and respective F2 progeny derived from backcrosses of F1 hybrid females to either parental strain (mixed genders). The black line indicates the mean cell number.

**(K)** Logarithm of odds (LOD) score across all four chromosomes for loci impacting Ir75b neuron numbers based on phenotyping data shown in **J**. Dashed horizontal lines mark *P* = 0.05; non-dashed horizontal lines mark *P* = 0.01.

**Figure S3: Odor-tracking behavior of wild-type strains.**

**(A)** Odor-tracking behavior towards noni juice in other wild-type strains of *D. sechellia* and *D. melanogaster* (see **Supplemental Table 2**), plotted as in Fig. 2B. Paired *t*-test, *** *P* < 0.001; ** *P* < 0.01; * *P* < 0.05; otherwise *P* > 0.05. *n* = 30 (*DmelBK, DmelrHR, Dsec19, Dsec31*) or 27 (*DmelOR, Dsec28*) animals.

**(B)** Odor-tracking behavior towards apple cider vinegar in wild-type *D. sechellia* strains. Paired *t*-test, *** *P* < 0.001; ** *P* < 0.01; * *P* < 0.05; otherwise *P* > 0.05. *n* = 30 animals each.

**Figure S4: Odor tracking in a wind tunnel.**

**(A)** Array of 12 tracking cameras above the wind tunnel used for 3D tracking of flies in the presence of a noni plume.

**(B)** Side view of wind tunnel system along with the acrylic port used to flux noni odor into the tunnel volume.

**(C)** Comparison of mean trajectory ground speeds for the three genotypes studied.

**(D)** Schematic of two plume models used to analyze data compared to the point cloud distribution of *D. sechellia* trajectories in the wind tunnel volume. A “straight” model (solid orange line) which is aligned in space with the odor port, or a “sloped” model (dashed orange line) for which the plume sinks by 5 cm from the odor port to the tunnel end.

**(E)** Radial distance calculations comparing mean radial distance of points from the plume centerline considering either the sloped or straight plume model (left two plots). The same data is also shown without the restriction to limit this analysis to 10 cm above or below the plume’s altitude (right two plots).

**(F)** Left, course direction distributions within a 3 cm radius of the plume centerline for either the sloped plume model (as in Fig. 2G) for the downwind and upwind halves of the wind tunnel, up to 5 cm downwind from the plume origin. Right, the same analysis assuming a straight plume model.

**Figure S5: Additional conditions of optogenetic stimulation.**

**(A)** Optic fiber-mediated illumination of one antenna in the tethered fly. The photo was taken from below the apparatus.

**(B)** Behavioral responses to optogenetic stimulation of Or22a/b OSNs in *D. melanogaster* using a *Gal4* knock-in line (see Fig. 3A for methodological details) Genotype: *D. melanogaster w;Or22a/b^Gal4^/UAS-CsChrimson-Venus*. Left, time course of Δ*WBA*. Right, quantification within each phase. Paired *t*-test, *P* > 0.05.

**(C)** Behavioral responses to optogenetic stimulation of effector control flies (*UAS-CsChrimson-Venus*) in *D. melanogaster* (left) and *D. sechellia* (right). Paired *t*-test*, P* > 0.05. Genotypes: *D. melanogaster w;+/UAS-CsChrimson-Venus, D. sechellia w;;;+/UAS-CsChrimson-Venus*.

**(D)** Behavioral responses to optogenetic stimulation of Or22a OSNs in *D. sechellia* using CsChrimson by stimulating the left antenna induces leftward turning behavior (resulting in reduced Δ*WBA* values compared to control flies). Paired *t*-test, ** *P* < 0.01; * *P* < 0.05; otherwise *P* > 0.05. Genotype as in Fig. 3A.

**Figure S6: SPARC2 allows genetic manipulation of a subset of Or22a OSNs.**

**(A)** Schematic illustrating the principle of SPARC2 (Isaacman-Beck et al., 2020). Top, ΦC31-mediated recombination leads to removal of a stop cassette flanked by attP sites. Co-expression of Gal4 in the same cell leads to transcriptional activation of a reporter gene. Middle, recombination with an alternative attP site does not lead to a functional read-out. Depending on the attP version present in the cassette, recombination efficiencies vary (D, dense; I, intermediate; S, sparse). Bottom, after recombination, a proportion of Gal4-expressing cells – in this study Or22a OSNs – will express the respective SPARC2 transgene.

**(B)** Left, immunofluorescence for GFP on whole-mount antennae from animals expressing different *SPARC2-X-GFP* versions in Or22a neurons. Right, quantification of the number of Or22a OSNs labelled by SPARC2 versions in *D. melanogaster*. Genotypes: *D. melanogaster nSyb-*Φ*C31/+;Or22a^Gal4^/UAS-SPARC2-X-GFP*. The SPARC2-D transgene labelled ∼50% of Or22a OSNs (compare with Fig. 1C) and was therefore used to generate transgenic constructs in *D. sechellia*.

**(C)** Optogenetic stimulation of Or22a OSNs in *D. sechellia* using a lower light intensity. Electrophysiological measurements confirm that this light intensity evokes a similar degree of OSN firing as in the SPARC2-D-CsChrimson experiment. Paired *t*-test. *\* P* < 0.05; otherwise *P* > 0.05. Electrophysiological data are re-plotted from Fig. 3A.

**(D)** Immunofluorescence for HA and nc82 on a whole-mount brain of *D. sechellia Or22a^Gal4^* transgenic flies expressing *UAS-SPARC2-D-TNT-HA-GeCO*, revealing selective strong expression in Or22a OSNs projecting to DM2. The white arrowheads point to a few cells (putative glia due to their apparent lack of processes) labelled in the central brain. Scale bar, 50 µm.

**Figure S7: Whole-cell patch clamp recordings from VM5d PNs.**

**(A)** Voltage trace (top) and peri-stimulus time histogram (PSTH, bottom) of VM5d PN responses to a dilution series of *2*-heptanone. Mean is shown for the voltage trace and mean ± SEM are shown for the PSTH. *n* = 2-5 animals.

**(B)** Start and end responses of VM5d PN to various dilutions of *2*-heptanone. Quantification at the start (first 50 ms), end (last 500 ms before odor offset) and decay magnitude (start - end) spike frequencies are shown. Student’s *t*-test. \**P* < 0.05. *n* = 2-5 animals (mean values and SEM for each concentration listed in Supplemental Table 1).

**Figure S8: Dose-response odor responses of OSNs and PNs in Or85b and Or22a pathways.**

**(A-B)** Representative odor-evoked calcium responses in the axon termini of Or85b OSNs (**A**) and dendrites of VM5d PNs (**B**) in *D. melanogaster* and *D. sechellia.* Far left, schematic of the OSN-PN populations under investigation; the population subject to imaging analysis in shown in a darker color. Left, representative confocal images of the raw fluorescence of GaMP6f showing the plane used for imaging with glomerular labels; corresponding solvent and odor stimulus-induced fluorescent changes are also shown. Middle panels, time courses in response to solvents and two odor dilutions (mean ± SEM Δ*F/F_0_*, bars indicate the timing of stimulus). Right, quantifications of peak Δ*F/F_0_*for a dilution series of each odor (Dunnett’s test, control: solvent, *** *P* < 0.001; * *P* < 0.01; * *P* < 0.05; otherwise *P* > 0.05, *n =* 7 (*D. melanogaster* OSNs), 8 (*D. sechellia* OSNs), 7 (*D. melanogaster* PNs), 6 (*D. sechellia* PNs) animals. Genotypes are indicated in Fig. 5G legend.

**(C-D)** Representative odor-evoked calcium responses in the axon termini of Or22a OSNs (**C**) and DM2 PNs (**D**) in *D. melanogaster* and *D. sechellia. n =* 10 animals each. Genotypes are indicated in Fig. 5G legend.

**(E)** Calcium imaging of DM2 PNs following reintroduction of *DmelOr22a* or *DsecOr22a* in Or22a OSNs in *D. melanogaster* lacking endogenous receptor expression. Left, time courses of odor responses. Middle, quantifications of peak Δ*F/F_0_* (Dunnett’s test, control: solvent, *** *P* < 0.001; * *P* < 0.01; * *P* < 0.05; otherwise *P* > 0.05,). *n =* 7 animals each. Right, normalized GCaMP6f fluorescence changes. Genotypes: *D. melanogaster w*;*Or22a/b^Gal4^*/*Or22a/b^Gal4^*;*UAS-DmelOr22a*,*VT033006-LexA*/*VT033006-LexA,LexAop-GCaMP6m* (*DmelOr22a* allele rescue), *w*;*Or22a/b^Gal4^*/*Or22a/b^Gal4^*;*UAS-DsecOr22a*,*VT033006-LexA*/*VT033006-LexA,LexAop-GCaMP6m* (*DsecOr22a* allele rescue).

**Figure S9: Odor response reliability.**

Odor response reliability of VM5d PNs. **(A-B)** Calcium responses of VM5d PNs in response to repeated stimulation with *2*-heptanone (concentration (Log) indicated on the figure) in *D. melanogaster* (**A**) and *D. sechellia* (**B**). Left, quantification of peak Δ*F/F_0_* from eight trials of stimulation. Dunnett’s test, compared to control (Trial 1). NS (*P* > 0.05). *n =* 8 animals each. Right, time courses of responses in trial 1 and 8 shown as mean ± SEM Δ*F/F_0_*. The odor stimulus (1 s) is shown with a black bar. **(C)** Quantification of mean, standard deviation and coefficient of variation of peak Δ*F/F_0_* across the eight stimulation trials. Welch’s *t*-test. NS (*P* > 0.05).

**Figure S10: *D. sechellia* VM5d PNs show persistent response to long odor stimulation**

Calcium responses of Or85b/VM5d neurons in response to long odor stimulation of *D. melanogaster* Or85b OSNs (**A**), *D. melanogaster* VM5d PNs (**B**), *D. sechellia* OSNs (**C**), and *D. sechellia* PNs (**D**). Left, time courses of the odor response (mean ± SEM Δ*F/F_0_*). Bars indicate the timing of long odor stimulation (10 s). Right, maximum Δ*F/F_0_*in the first and last 2.5 s time windows are compared. Paired *t*-test. ** *P* < 0.01; ** *P* < 0.05. *n =* 7 (OSNs) or 8 (PNs) animals for both species. Genotypes are indicated in the Fig. 5G legend.

**Figure S11: Odor response sensitivity in control glomeruli.**

Odor-evoked dose-dependent calcium responses in control glomeruli of *D. melanogaster* VA2 (Or92a) PNs (**A**), *D. sechellia* VA2 PNs (**B**)*, D. melanogaster* DM4 (Or59b) PNs (**C**), and *D. sechellia* DM4 PNs (**D**). Left, time courses in response to solvents and specific odor dilutions (mean ± SEM Δ*F/F_0_*, bars indicate the timing of stimuli). Right, quantifications of peak Δ*F/F_0_* (Dunnett’s test, control: solvent, *** *P* < 0.001; * *P* < 0.01; * *P* < 0.05; otherwise *P* > 0.05,). *n =* 7 each. The genotype for the VA2 and DM4 PN calcium imaging is the same as for DM2 PN imaging shown in Fig. 5G.

**Figure S12: Pharmacological manipulations of lateral inhibition.**

**(A,B)** Odor pulse responses of VM5d PNs following application of a GABA_A_ (**A**, Picrotoxin) and GABA_B_ (**B**, CGP54626) antagonist. PN responses in normal AHL saline (left) or containing 100 µM picrotoxin or 50 µM CGP54626 (middle). The ratio between 10^th^ and 1^st^ odor pulses are shown on the far right. Paired *t*-test. ** *P* < 0.01, * *P* < 0.05, NS *P* > 0.05. *n =* 7 animals each. Genotypes are indicated in Fig. 5G legend.

**c**, Raw GCaMP fluorescence intensity in VM5d PNs in *D. melanogaster* (left) and *D. sechellia* (right) following GABA antagonist application. Mean fluorescence before (4-5 s; as "Baseline") and during (5-9 s; as “Odor”) the stimulus was quantified. Paired *t*-test, ** *P* < 0.01, * *P* < 0.05, NS *P* > 0.05. *n =* 8 animals each. Genotype as in Fig. 5G.

**Figure S13: Variation in neuron numbers within and across species.**

**(A)** Correlation between OSN number and synapse numbers across glomeruli from *D. melanogaster* OSN quantification (Grabe et al., 2016) and connectome (Schlegel et al., 2021) data. OSN outputs were quantified separately for connections to PNs (left), LNs (middle), and OSNs (right). *r* and *p* values are calculated by Pearson’s correlation analysis.

**(B)** Variation in basiconic sensilla neuron numbers in the *D. melanogaster* species subgroup. Numbers of Or42b, Or59b and Or22a/(b) neurons (housed in ab1, ab2 and ab3 sensilla, respectively) based on RNA FISH on whole-mount antennae across the *D. melanogaster* species subgroup; Ma, million years ago (n = 7-14, females). For strain details see **Supplemental Table 2**.

## Supplemental Tables

**Supplemental Table 1:** Spike counts for electrophysiological experiments.

**Supplemental Table 2:** Wild-type and transgenic lines used and generated in this study.

**Supplemental Table 3:** Oligonucleotides used to generate single sgRNA expression vectors and *in situ* probe templates.

**Supplemental Table 4:** Oligonucleotides used to generate multi-sgRNA expression vectors.

