## Supplementary Figures for "Sensory neuron population expansion enhances odor tracking without sensitizing projection neurons"

### Figure S1

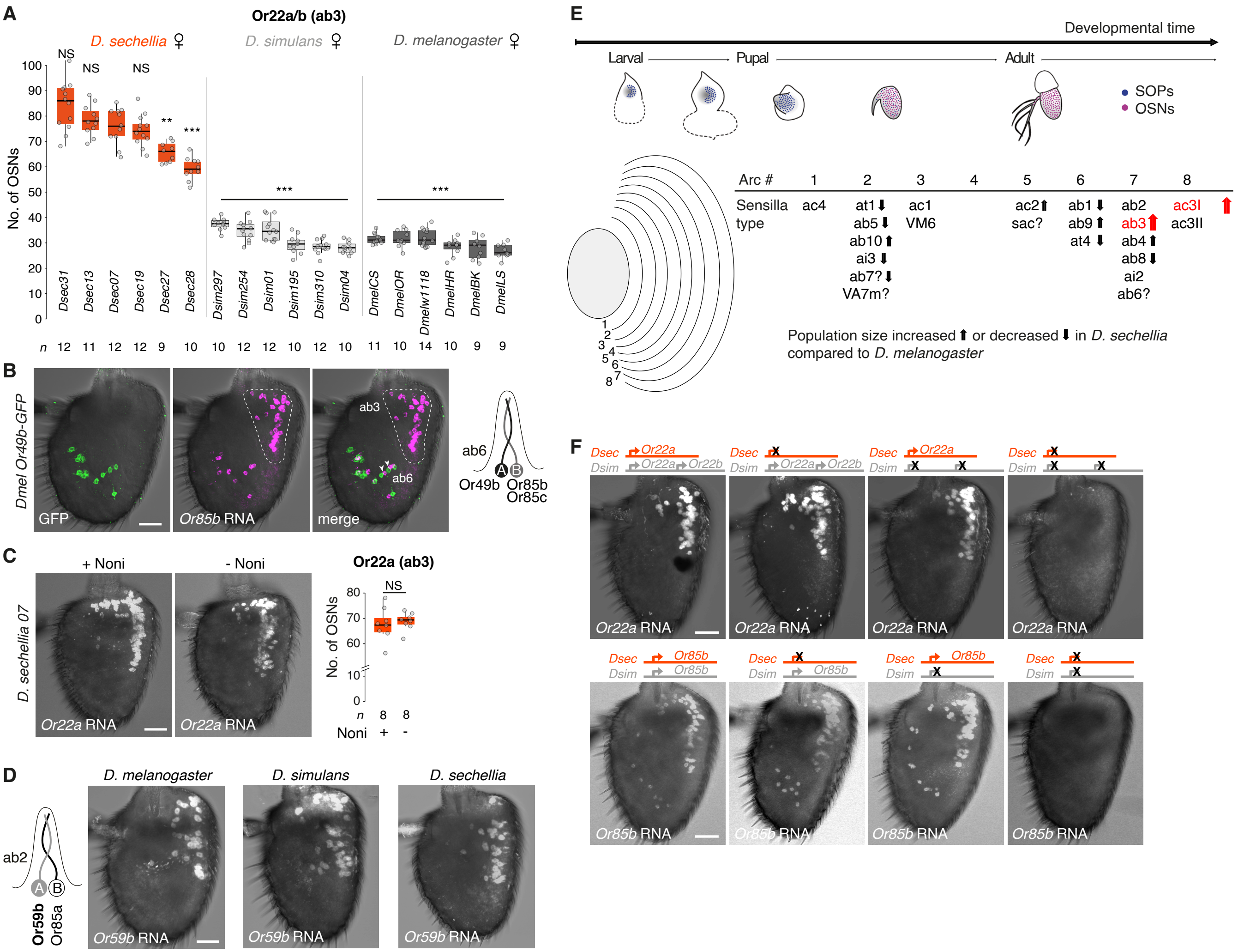
A
Or22a/b (ab3)
D. sechellia ♀
D. simulans ♀
D. melanogaster ♀
No. of OSNs
NS
NS
NS
**
***
***
***
Dsec31 Dsec13 Dsec07 Dsec19 Dsec27 Dsec28
Dsim297 Dsim254 Dsim01 Dsim195 Dsim310 Dsim04
DmelCS DmelOR Dmelw1118 DmelHR DmelBK DmelLS
n 12 11 12 12 9 10 10 12 12 10 12 10 11 10 14 10 9 9
B
Dmel Or49b-GFP
GFP
Or85b RNA
merge
ab3
ab6
ab6
Or49b Or85b Or85c
C
D. sechellia 07
+ Noni
- Noni
Or22a RNA
Or22a RNA
Or22a (ab3)
NS
No. of OSNs
n 8 8
Noni + -
D
ab2
Or59b Or85a
D. melanogaster
D. simulans
D. sechellia
Or59b RNA
Or59b RNA
Or59b RNA
E
Developmental time
Larval
Pupal
Adult
SOPs
OSNs
Arc # 1 2 3 4 5 6 7 8
Sensilla type
ac4
at1⬇ ab5⬇ ab10⬆ ai3⬇ ab7?⬇ VA7m?
ac1 VM6
ac2⬆ sac?
ab1⬇ ab9⬆ at4⬇
ab2 ab3⬆ ab4⬆ ab8⬇ ai2 ab6?
ac3I⬆ ac3II
Population size increased ⬆ or decreased ⬇ in D. sechellia compared to D. melanogaster
F
Dsec Or22a
Dsim Or22a Or22b
Or22a RNA
Dsec Or85b
Dsim Or85b
Or85b RNA

**Figure S2**

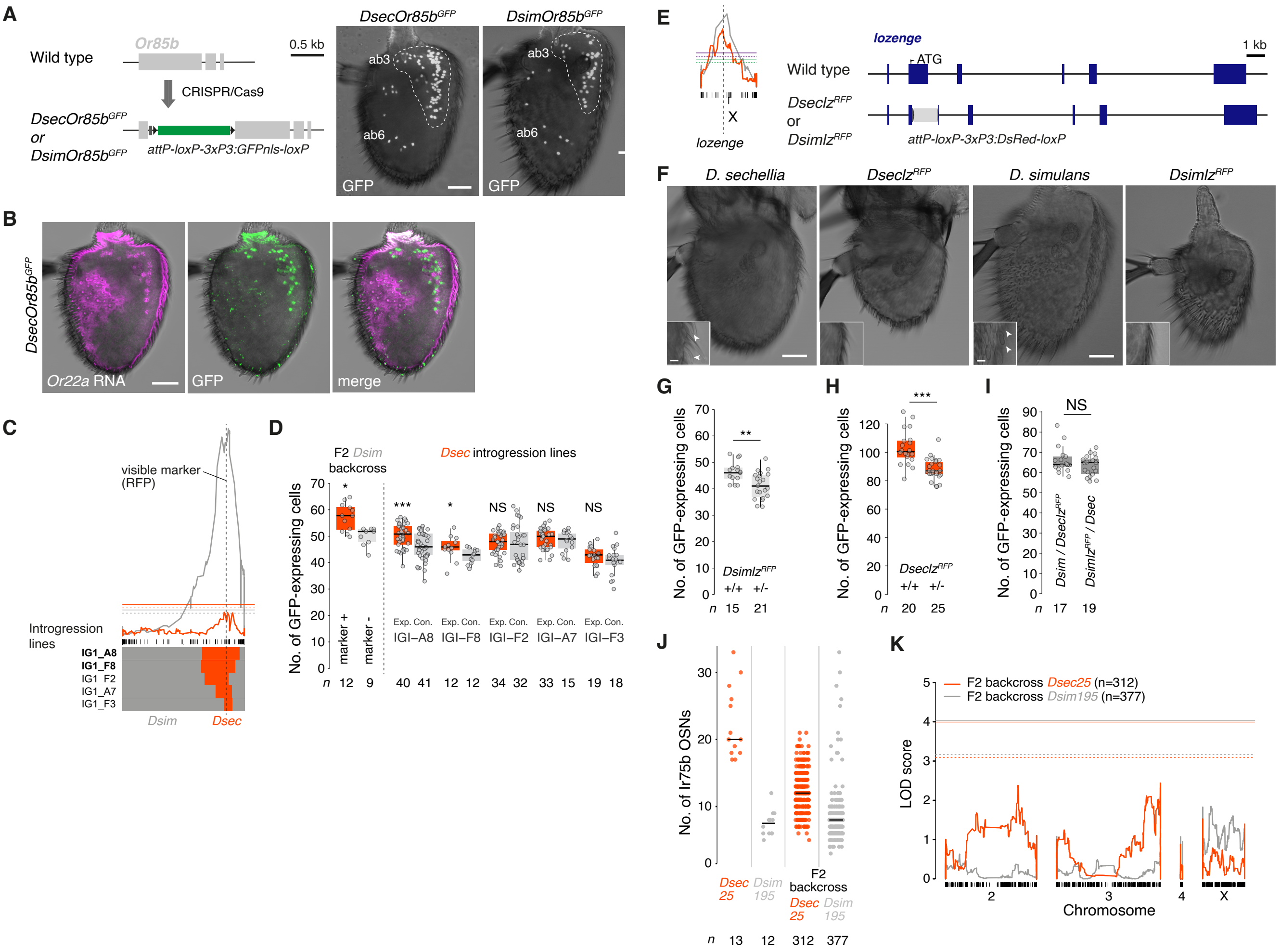
A
Wild type
Or85b
0.5 kb
CRISPR/Cas9
DsecOr85bGFP
or
DsimOr85bGFP
attP-loxP-3xP3:GFPnls-loxP
DsecOr85bGFP
DsimOr85bGFP
ab3
ab6
GFP
B
DsecOr85bGFP
Or22a RNA
GFP
merge
C
visible marker
(RFP)
Introgression
lines
IG1_A8
IG1_F8
IG1_F2
IG1_A7
IG1_F3
Dsim
Dsec
D
F2 Dsim
backcross
Dsec introgression lines
No. of GFP-expressing cells
marker +
marker -
Exp. Con.
IGI–A8
IGI–F8
IGI–F2
IGI–A7
IGI–F3
NS
n 12 9 40 41 12 12 34 32 33 15 19 18
E
lozenge
1 kb
ATG
Wild type
DseclzRFP
or
DsimlzRFP
attP-loxP-3xP3:DsRed-loxP
X
F
D. sechellia
DseclzRFP
D. simulans
DsimlzRFP
G
No. of GFP-expressing cells
DsimlzRFP
+/+ +/-
n 15 21
H
DseclzRFP
+/+ +/-
n 20 25
I
NS
Dsim / DseclzRFP
DsimlzRFP / Dsec
n 17 19
J
No. of Ir75b OSNs
Dsec 25
Dsim 195
F2 backcross
Dsec 25
Dsim 195
n 13 12 312 377
K
F2 backcross Dsec25 (n=312)
F2 backcross Dsim195 (n=377)
LOD score
Chromosome
2
3
4
X

### Figure S3

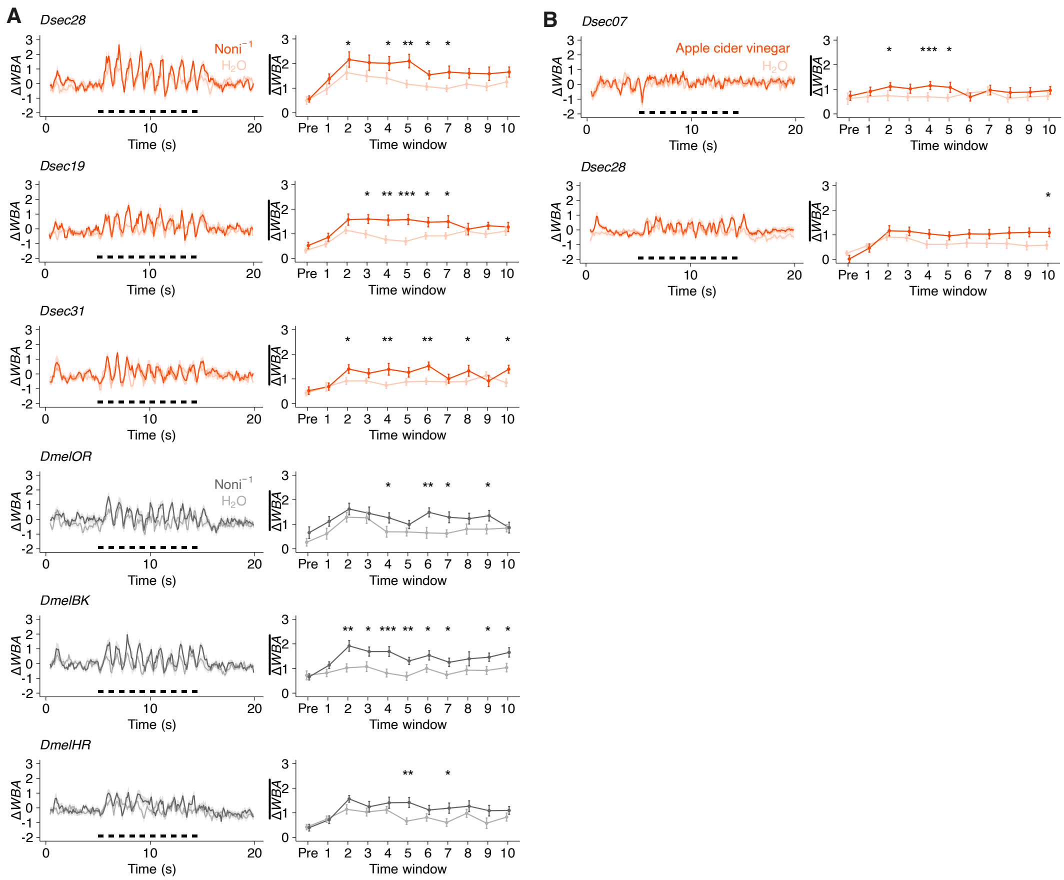
A
Dsec28
Noni$^{-1}$
$H_2O$
Dsec19
Dsec31
DmelOR
DmelBK
DmelHR
ΔWBA
Time (s)
Time window
B
Dsec07
Apple cider vinegar
Dsec28

**Figure S4**

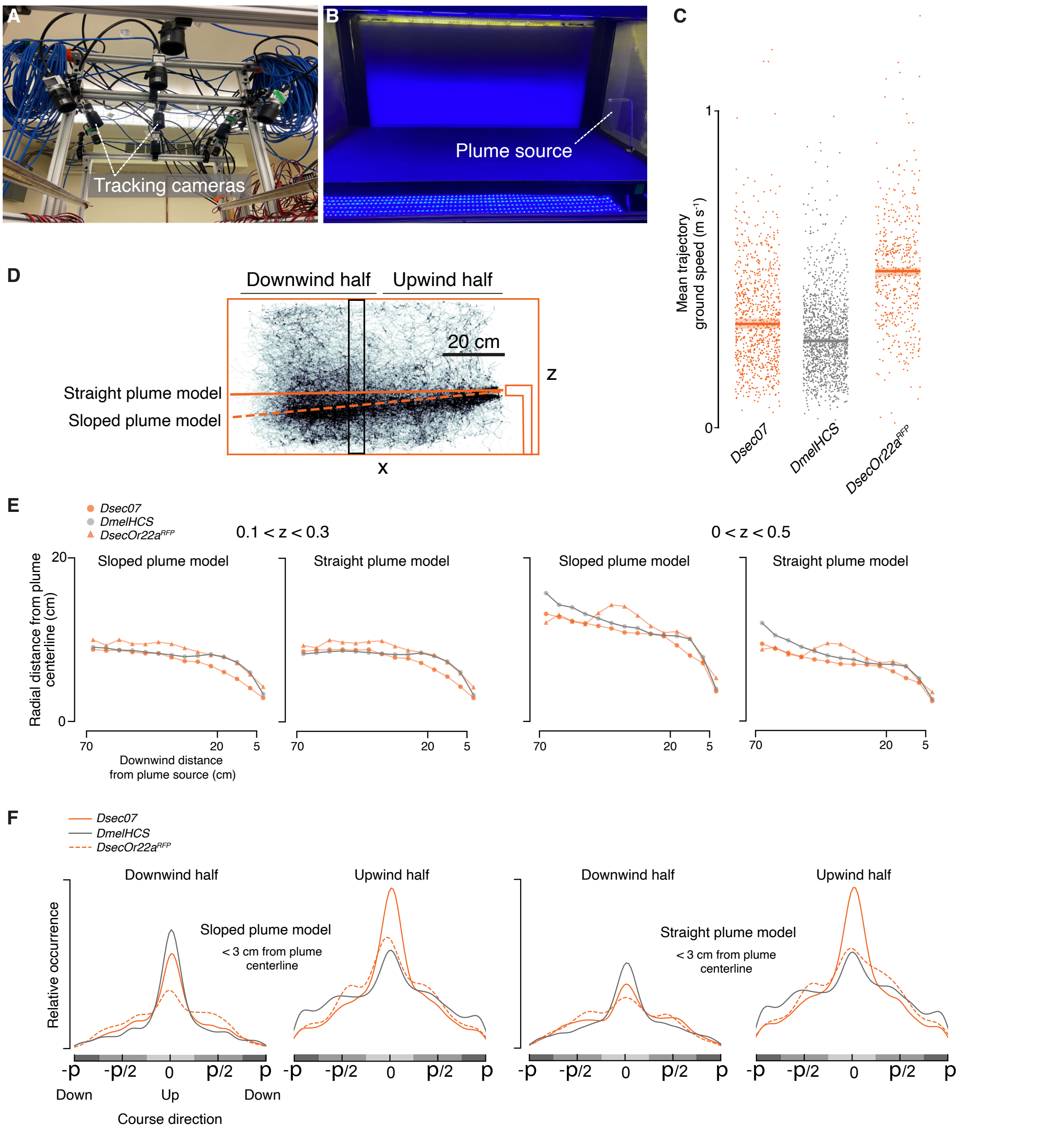

A
Tracking cameras
B
Plume source
C
1
0
Mean trajectory ground speed (m s-1)
Dsec07
DmelHCS
DsecOr22a^RFP
D
Downwind half
Upwind half
20 cm
z
x
Straight plume model
Sloped plume model
E
Dsec07
DmelHCS
DsecOr22a^RFP
0.1 < z < 0.3
0 < z < 0.5
Sloped plume model
Straight plume model
Sloped plume model
Straight plume model
20
0
Radial distance from plume centerline (cm)
70
20
5
Downwind distance from plume source (cm)
F
Dsec07
DmelHCS
DsecOr22a^RFP
Downwind half
Upwind half
Downwind half
Upwind half
Sloped plume model
< 3 cm from plume centerline
Straight plume model
< 3 cm from plume centerline
Relative occurrence
-p
-p/2
0
p/2
p
Down
Up
Down
Course direction

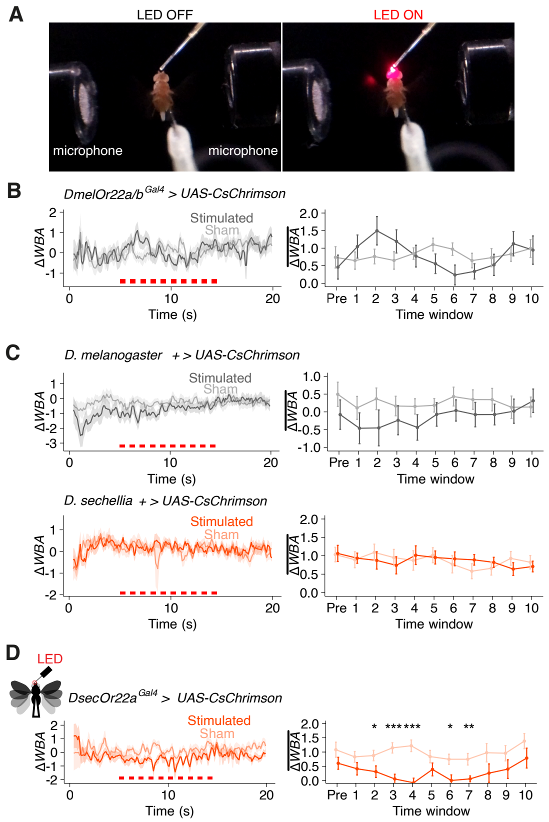
A
LED OFF
LED ON
microphone
microphone
B
DmelOr22a/b^Gal4 > UAS-CsChrimson
Stimulated
Sham
ΔWBA
Time (s)
Time window
C
D. melanogaster + > UAS-CsChrimson
D. sechellia + > UAS-CsChrimson
D
LED
DsecOr22a^Gal4 > UAS-CsChrimson

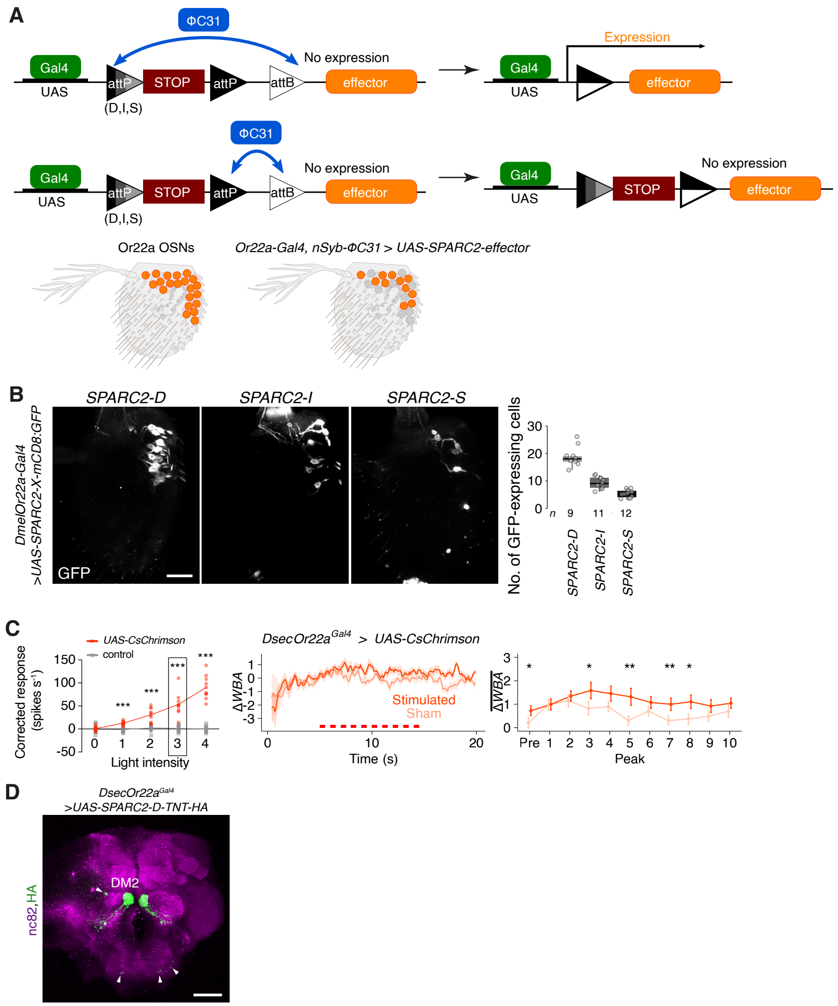
A
ΦC31
Gal4
UAS
attP
STOP
attP
attB
(D,I,S)
No expression
effector
Expression
Or22a OSNs
Or22a-Gal4, nSyb-ΦC31 > UAS-SPARC2-effector
B
SPARC2-D
SPARC2-I
SPARC2-S
DmelOr22a-Gal4 >UAS-SPARC2-X-mCD8:GFP
GFP
No. of GFP-expressing cells
n 9 11 12
C
UAS-CsChrimson
control
Corrected response (spikes s⁻¹)
Light intensity
DsecOr22a^Gal4 > UAS-CsChrimson
ΔWBA
Stimulated
Sham
Time (s)
Peak
D
DsecOr22a^Gal4 >UAS-SPARC2-D-TNT-HA
nc82,HA
DM2

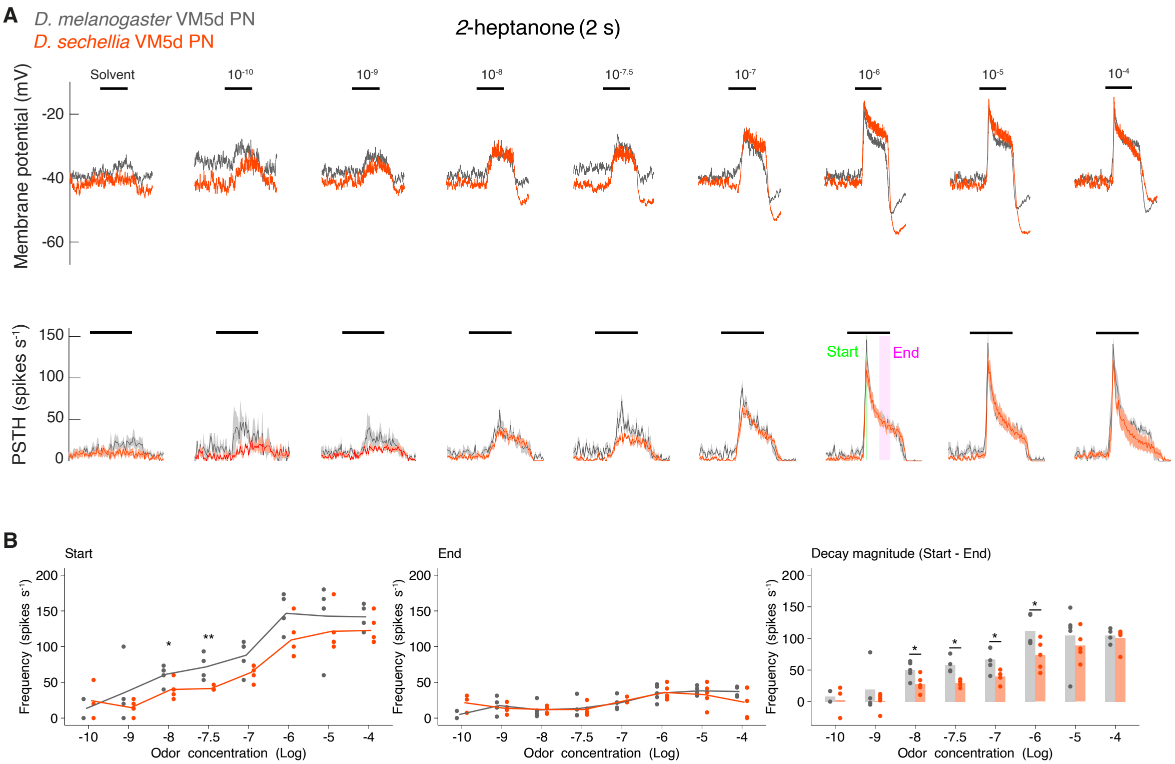
A
D. melanogaster VM5d PN
D. sechellia VM5d PN
2-heptanone (2 s)
Solvent
10-10
10-9
10-8
10-7.5
10-7
10-6
10-5
10-4
Membrane potential (mV)
-20
-40
-60
PSTH (spikes s-1)
150
100
50
0
Start
End
B
Start
End
Decay magnitude (Start - End)
Frequency (spikes s-1)
200
150
100
50
0
-10
-9
-8
-7.5
-7
-6
-5
-4
Odor concentration (Log)
*
**

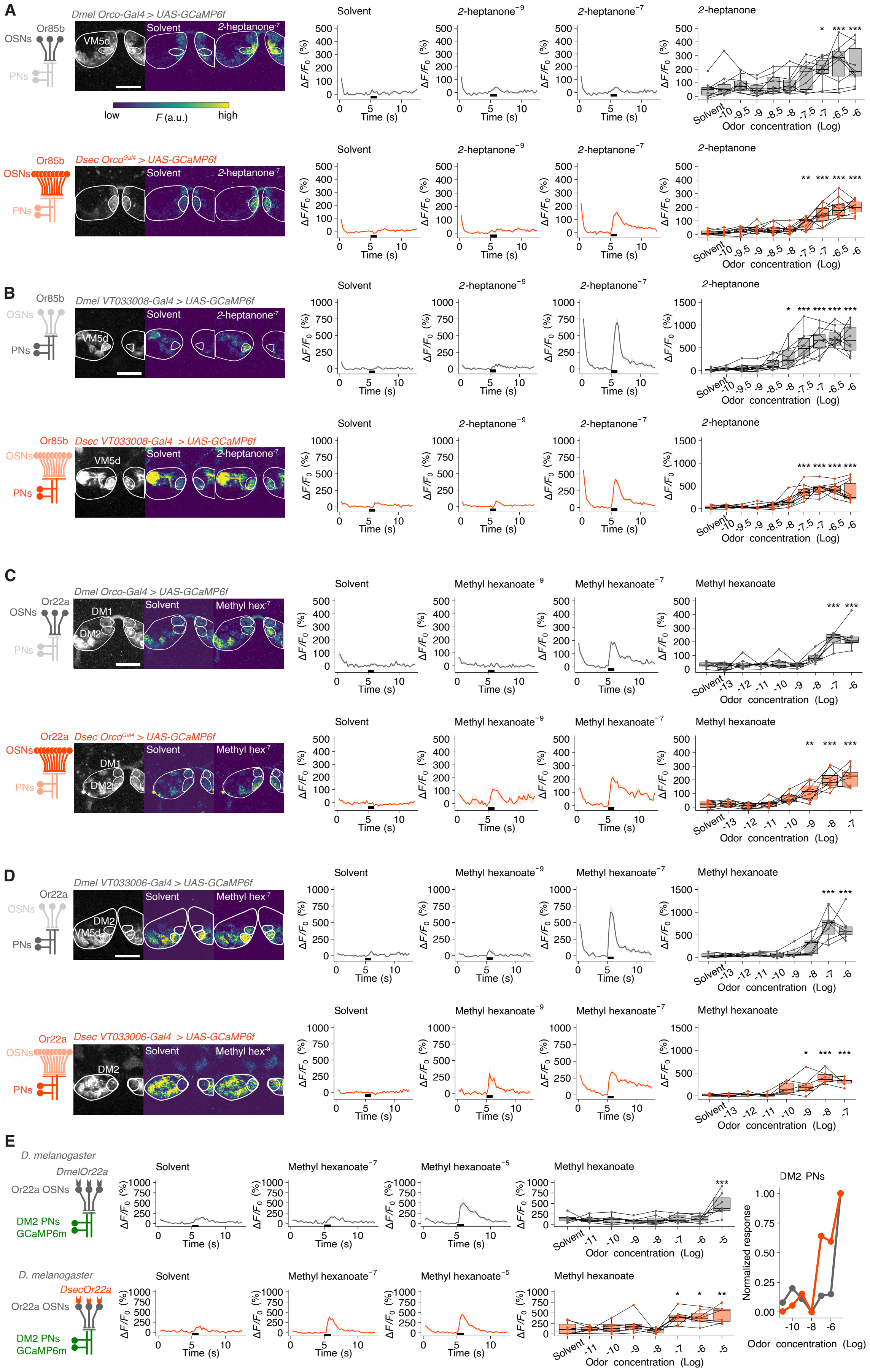
A
Or85b
OSNs
PNs
Dmel Orco-Gal4 > UAS-GCaMP6f
VM5d
Solvent
2-heptanone-7
low
F (a.u.)
high
Solvent
2-heptanone-9
2-heptanone-7
2-heptanone
ΔF/F0 (%)
Time (s)
Odor concentration (Log)
Dsec Orco-Gal4 > UAS-GCaMP6f
B
Dmel VT033008-Gal4 > UAS-GCaMP6f
Dsec VT033008-Gal4 > UAS-GCaMP6f
C
Or22a
Dmel Orco-Gal4 > UAS-GCaMP6f
DM1
DM2
Methyl hex-7
Methyl hexanoate-9
Methyl hexanoate-7
Methyl hexanoate
Dsec Orco-Gal4 > UAS-GCaMP6f
D
Dmel VT033006-Gal4 > UAS-GCaMP6f
Dsec VT033006-Gal4 > UAS-GCaMP6f
Methyl hex-9
E
D. melanogaster
DmelOr22a
Or22a OSNs
DM2 PNs
GCaMP6m
DsecOr22a
Methyl hexanoate-5
DM2 PNs
Normalized response

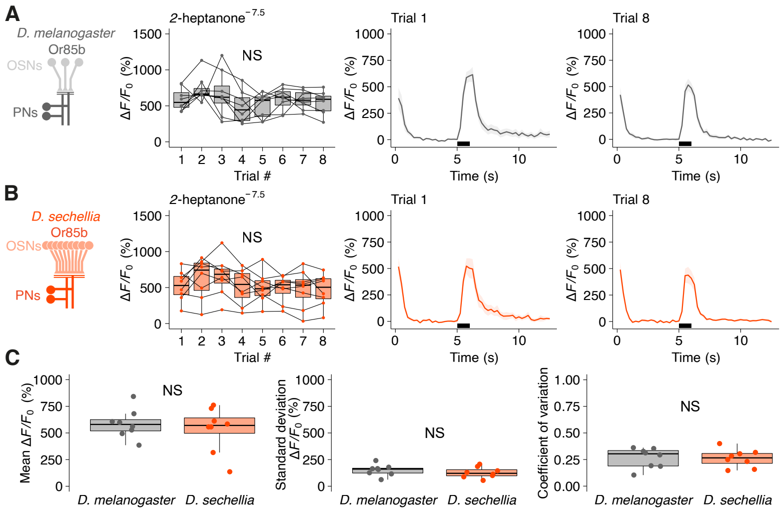
A
D. melanogaster
Or85b
OSNs
PNs
2-heptanone$^{-7.5}$
NS
Trial 1
Trial 8
ΔF/F0 (%)
Trial #
Time (s)
B
D. sechellia
Or85b
OSNs
PNs
2-heptanone$^{-7.5}$
NS
Trial 1
Trial 8
ΔF/F0 (%)
Trial #
Time (s)
C
Mean ΔF/F0 (%)
Standard deviation ΔF/F0 (%)
Coefficient of variation
NS
D. melanogaster
D. sechellia

### Figure S10

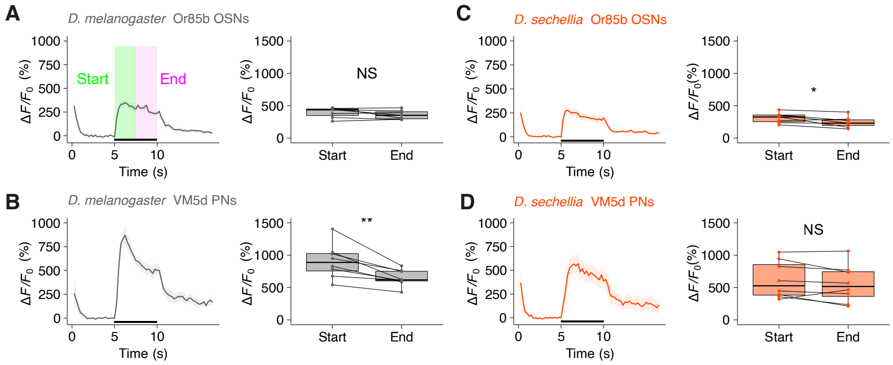

A
D. melanogaster Or85b OSNs
Start
End
ΔF/F0 (%)
Time (s)
NS
B
D. melanogaster VM5d PNs
**
C
D. sechellia Or85b OSNs
*
D
D. sechellia VM5d PNs
NS

### Figure S11

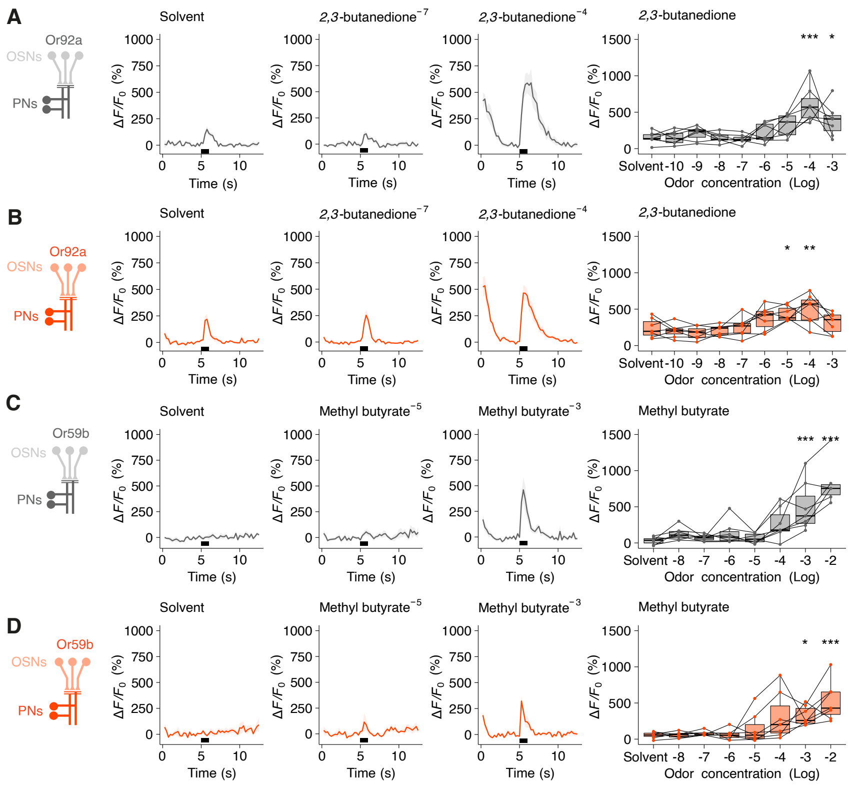

A
Or92a
OSNs
PNs
Solvent
2,3-butanedione$^{-7}$
2,3-butanedione$^{-4}$
2,3-butanedione
ΔF/F$_0$ (%)
Time (s)
Odor concentration (Log)
Solvent -10 -9 -8 -7 -6 -5 -4 -3
*** *
B
Or92a
OSNs
PNs
Solvent
2,3-butanedione$^{-7}$
2,3-butanedione$^{-4}$
2,3-butanedione
* **
C
Or59b
OSNs
PNs
Solvent
Methyl butyrate$^{-5}$
Methyl butyrate$^{-3}$
Methyl butyrate
Solvent -8 -7 -6 -5 -4 -3 -2
*** ***
D
Or59b
OSNs
PNs
Solvent
Methyl butyrate$^{-5}$
Methyl butyrate$^{-3}$
Methyl butyrate
* ***

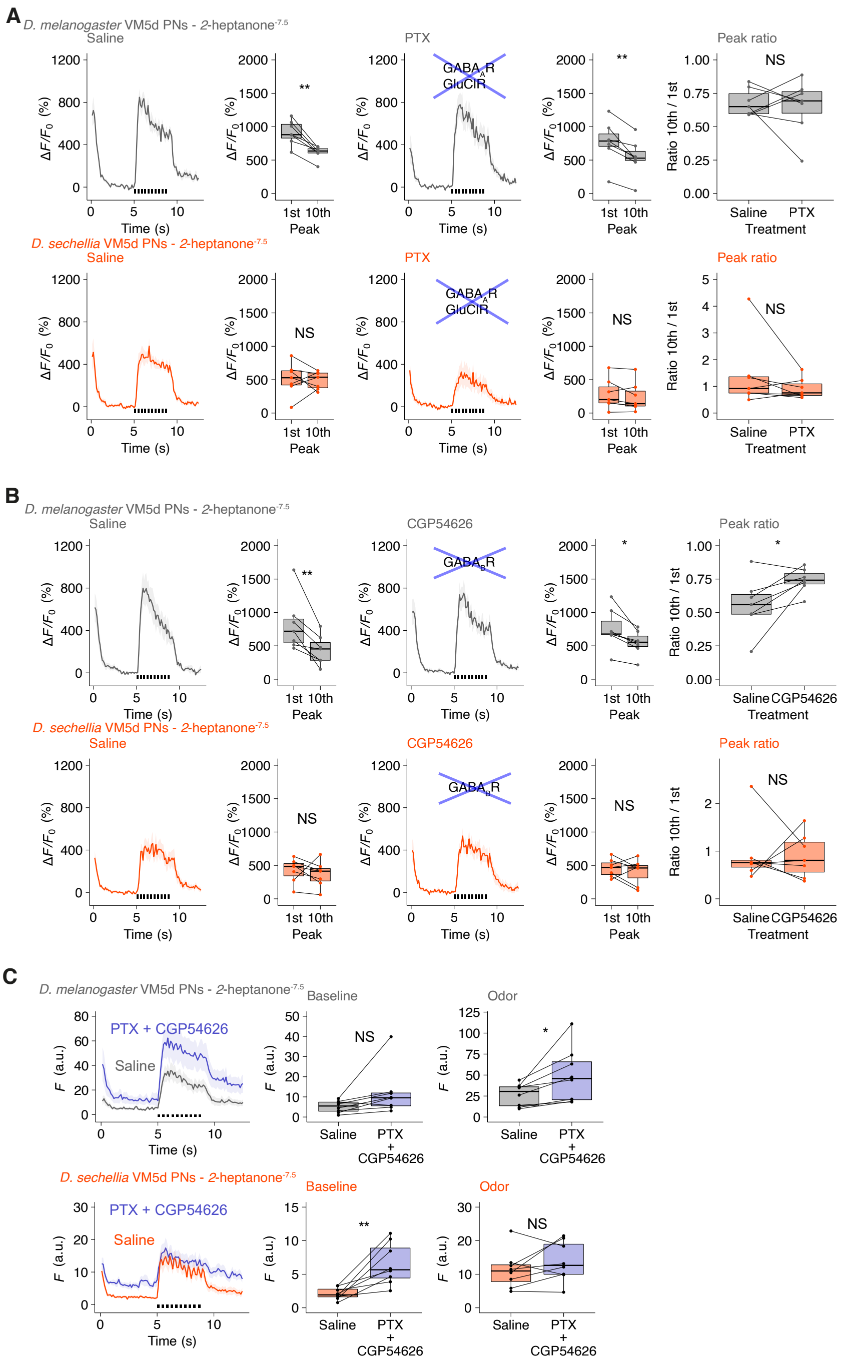
A
D. melanogaster VM5d PNs - 2-heptanone-7.5
Saline
PTX
GABAA R
GluClR
Peak ratio
NS
**
Ratio 10th / 1st
Saline PTX
Treatment
1st 10th
Peak
Time (s)
ΔF/F0 (%)
D. sechellia VM5d PNs - 2-heptanone-7.5
B
CGP54626
GABAB R
*
C
PTX + CGP54626
Saline
Baseline
Odor
F (a.u.)
PTX + CGP54626

**Figure S13**

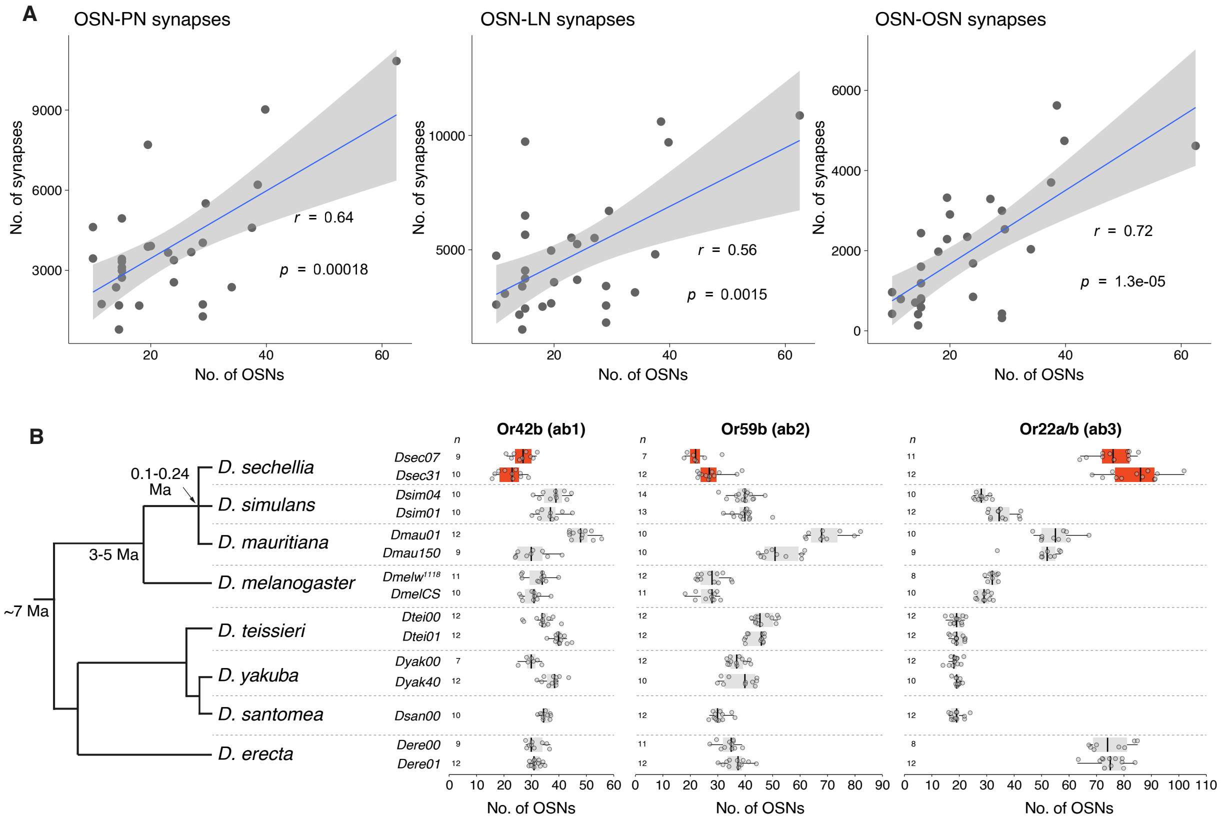

A
OSN-PN synapses
OSN-LN synapses
OSN-OSN synapses
No. of synapses
No. of OSNs
r = 0.64
p = 0.00018
r = 0.56
p = 0.0015
r = 0.72
p = 1.3e-05
B
~7 Ma
3-5 Ma
0.1-0.24 Ma
D. sechellia
D. simulans
D. mauritiana
D. melanogaster
D. teissieri
D. yakuba
D. santomea
D. erecta
Or42b (ab1)
Or59b (ab2)
Or22a/b (ab3)
n
Dsec07
Dsec31
Dsim04
Dsim01
Dmau01
Dmau150
Dmelw1118
DmelCS
Dtei00
Dtei01
Dyak00
Dyak40
Dsan00
Dere00
Dere01
No. of OSNs
