## Supplementary material for "Sensory neuron population expansion enhances odor tracking without sensitizing projection neurons": Table S2-4

**Supplemental Table 2.** **Wild-type and transgenic lines used and generated in this study.**

| **Stock name** | **Donor plasmid** | **Parental strain** | **Species** | **Method or Reference** | **Figure** |
| --- | --- | --- | --- | --- | --- |
| *Dsec07* |  | *Drosophila* Species Stock Center [DSSC] 14021-0248.07 | *D. sechellia* |  | Figure 1,  Figure 2,  Suppl. Fig. 1,  Suppl. Fig. 2,  Suppl. Fig. 3,  Suppl. Fig. 4,  Suppl. Fig. 13 |
| *Dsec31* |  | DSSC 14021-0248.31 | *D. sechellia* |  | Suppl. Fig. 1,  Suppl. Fig. 3,  Suppl. Fig. 13 |
| *Dsec19* |  | DSSC 14021-0248.19 | *D. sechellia* |  | Suppl. Fig. 1,  Suppl. Fig. 3 |
| *Dsec13* |  | DSSC 14021-0248.13 | *D. sechellia* |  | Suppl. Fig. 1 |
| *Dsec27* |  | DSSC 14021-0248.27 | *D. sechellia* |  | Suppl. Fig. 1 |
| *Dsec28* |  | DSSC 14021-0248.28 | *D. sechellia* |  | Suppl. Fig. 1,  Suppl. Fig. 3 |
| *Dsec25* |  | DSSC 14021-0248.25 | *D. sechellia* |  | Suppl. Fig. 2 |
| *Dsim297* |  | DSSC 14021-0251.297 | *D. simulans* |  | Suppl. Fig. 1 |
| *Dsim254* |  | DSSC 14021-0251.254 | *D. simulans* |  | Suppl. Fig. 1 |
| *Dsim01* |  | DSSC 14021-0251.001 | *D. simulans* |  | Suppl. Fig. 1,  Suppl. Fig. 13 |
| *Dsim195* |  | DSSC 14021-0251.195 | *D. simulans* |  | Suppl. Fig. 1  Suppl. Fig. 2 |
| *Dsim310* |  | DSSC 14021-0251.310 | *D. simulans* |  | Suppl. Fig. 1 |
| *Dsim04* |  | DSSC 14021-0251.004 | *D. simulans* |  | Figure 1,  Suppl. Fig. 1,  Suppl. Fig. 2,  Suppl. Fig. 13 |
| *Dsim03* |  | DSSC 14021-0251.003 | *D. simulans* |  | Suppl. Fig. 2 |
| *DmelCS* |  | *D. melanogaster* *Canton-S* | *D. melanogaster* |  | Figure 1,  Figure 2,  Suppl. Fig. 1, Suppl. Fig. 13 |
| *DmelOR* |  | *Dmel Oregon-R* | *D. melanogaster* |  | Suppl. Fig. 1,  Suppl. Fig. 3 |
| *Dmelw^1118^* |  | *Dmel w^1118^* | *D. melanogaster* |  | Suppl. Fig. 1,  Suppl. Fig. 13 |
| *DmelHR* |  | *Dmel Hikone-R* | *D. melanogaster* |  | Suppl. Fig. 1,  Suppl. Fig. 3 |
| *DmelBK* |  | *Dmel Berlin-K* | *D. melanogaster* |  | Suppl. Fig. 1,  Suppl. Fig. 3 |
| *DmelLS* |  | *Dmel Lausanne-S* | *D. melanogaster* |  | Suppl. Fig. 1 |
| *DmelHCS* |  | *Dmel Heisenberg CS* | *D. melanogaster* |  | Figure 2,  Suppl. Fig. 4 |
| *Dmau01* |  | DSSC 14021-0241.01 | *D. mauritiana* |  | Suppl. Fig. 13 |
| *Dmau150* |  | DSSC 14021-0241.150 | *D. mauritiana* |  | Suppl. Fig. 13 |
| *Dtei00* |  | DSSC 14021-0257.00 | *D. teissieri* |  | Suppl. Fig. 13 |
| *Dtei01* |  | DSSC 14021-0257.01 | *D. teissieri* |  | Suppl. Fig. 13 |
| *Dyak00* |  | DSSC 14021-0261.00 | *D. yakuba* |  | Suppl. Fig. 13 |
| *Dyak40* |  | DSSC 14021-0261.40 | *D. yakuba* |  | Suppl. Fig. 13 |
| *Dsan00* |  | DSSC 14021-0271.00 | *D. santomea* |  | Suppl. Fig. 13 |
| *Dere00* |  | DSSC 14021-0224.00 | *D. erecta* |  | Suppl. Fig. 13 |
| *Dere01* |  | DSSC 14021-0224.01 | *D. erecta* |  | Suppl. Fig. 13 |
| *DsecOr22a^RFP^* |  |  | *D. sechellia* | (Auer et al., 2020) | Figure 1,  Figure 2,  Suppl. Fig. 1,  Suppl. Fig. 4 |
| *DsecOr85c/b^RFP^* |  |  | *D. sechellia* | (Auer et al., 2020) | Figure 2 |
| *DsecIr75b^RFP^* |  |  | *D. sechellia* | (Auer et al., 2020) | Figure 2 |
| *DsecOr35a^RFP^* |  |  | *D. sechellia* | (Auer et al., 2020) | Figure 2 |
| *DsimOr85b^GFP^* | *DsimOr85b,3xP3-Stinger* | DSSC 14021-0251.003 | *D. simulans* | CRISPR knock-in, this study | Suppl. Fig. 2 |
| *DsimOr85b^GFP^* | *DsimOr85b,3xP3-Stinger* | DSSC 14021-0251.004 | *D. simulans* | CRISPR knock-in, this study | Figure 1,  Suppl. Fig. 1,  Suppl. Fig. 2 |
| *DsecOr85b^GFP^* | *DsecOr85b,3xP3-Stinger,* | DSSC 14021-0248.07 | *D. sechellia* | (Auer et al., 2020) | Figure 1,  Suppl. Fig. 1,  Suppl. Fig. 2 |
| *Dsec pBAC(UAS-CsChrimson-Venus)* | *pBAC(UAS-CsChrimson-Venus-3xP3-DsRed)* |  | *D. sechellia* | *pBAC* integration, this study | Figure 3,  Suppl. Fig. 2,  Suppl. Fig. 5 |
| *Dseclz^RFP^* | *pHD-3xP3-DsRed-Dseclz* | *Dsecnanos-Cas9* | *D. sechellia* | CRISPR knock-in, this study | Suppl. Fig. 2 |
| *Dsimlz^RFP^* | *pHD-3xP3-DsRed-Dsimlz* | DSSC 14021-0251.004 | *D. simulans* | CRISPR knock-in, this study | Suppl. Fig. 2 |
| *DsimOr22a/b^RFP^* |  |  | *D. simulans* | (Auer et al., 2020) | Figure 1,  Suppl. Fig. 1 |
| *Dmel Or49b-GFP* |  | Bloomington *Drosophila* Stock Center [BDSC]_52653 | *D. melanogaster* | (Couto et al., 2005) | Suppl. Fig. 1 |
| *Dsec UAS-GCaMP6f* |  |  | *D. sechellia* | (Auer et al., 2020) | Figure 4,  Figure 5,  Figure 6,  Suppl. Fig. 8,  Suppl. Fig. 9,  Suppl. Fig. 10,  Suppl. Fig. 11,  Suppl. Fig. 12 |
| *Dsec VT033006-Gal4* | *VT033006-Gal4 (from* *(Tirian and Dickson, 2017))* | *Dsecwhite* (Auer et al., 2020) | *D. sechellia* | attB/P integration, this study | Figure 4,  Figure 5,  Figure 6,  Suppl. Fig. 8,  Suppl. Fig. 9,  Suppl. Fig. 10,  Suppl. Fig. 11,  Suppl. Fig. 12 |
| *Dsec VT033008-Gal4* | *VT033008-Gal4 (from(Tirian and Dickson, 2017))* | *Dsecwhite (Auer et al., 2020)* | *D. sechellia* | attB/P integration, this study | Figure 4 |
| *Dsec UAS-C3PA-GFP* |  |  | *D. sechellia* | (Auer et al., 2020) | Figure 4 |
| *Dsec UAS-D*α*7-GFP* | *p(UAS-D*α*7-GFP) (gift from S. Sigrist* (Leiss et al., 2009)) | DSSC 14021-0248.30 | *D. sechellia* | P-element transgenesis, this study | Figure 4 |
| *Dmel VT033006-Gal4* |  | *Vienna Drosophila Resource Center* [VDRC] ID 202281 | *D. melanogaster* | (Tirian and Dickson, 2017) | Figure 4,  Figure 5,  Figure 6,  Suppl. Fig. 8,  Suppl. Fig. ,  9,  Suppl. Fig. 11,  Suppl. Fig. 12 |
| *Dmel VT033008-Gal4* |  | VDRC_ID 200242 | *D. melanogaster* | (Tirian and Dickson, 2017) | Figure 4 |
| *Dmel UAS-C3PA-GFP* |  |  | *D. melanogaster* | (Ellis et al., 2023) | Figure 4 |
| *Dmel UAS-D*α*7-GFP* |  |  | *D. melanogaster* | (Leiss et al., 2009) | Figure 4 |
| *Dsec VM5d-Gal4* | *GMR_86C10-GAL4 (Jenett et al., 2012)* | *Dsec-attP40* (T.O.A. *unpublished*) | *D. sechellia* | attB/P integration | Figure 4,  Figure 5,  Suppl. Fig. 7 |
| *Dsec UAS-myrGFP* | *pUAS-myrGFP, QUAS-mtdTomato(3xHA)* (Talay et al., 2017) | *Dsec-attP40* (T.O.A. *unpublished*) | *D. sechellia* | attB/P integration | Figure 4,  Figure 5,  Suppl. Fig. 7 |
| *ODmel Vm5d-Gal4* |  | BDSC_46820 | *D. melanogaster* |  | Figure 4,  Figure 5,  Suppl. Fig. 7 |
| *Dmel UAS-GFP* |  | BDSC_52262 | *D. melanogaster* |  | Figure 4,  Figure 5,  Suppl. Fig. 7 |
| *Dmel UAS-DsecOr22a* |  |  | *D. melanogaster* | (Auer et al., 2020) | Suppl. Fig. 8 |
| *Dmel UAS-DmelOr22a* |  |  | *D. melanogaster* | (Auer et al., 2020) | Suppl. Fig. 8 |
| *Dmel VT33006-LexA* | *p(VT033006-LexA)* | *D. melanogaster* attP2 | *D. melanogaster* | attB/P integration, this study | Suppl. Fig. 8 |
| *Dmel LexAop-GCaMP6m* |  | BCSC_44588 | *D. melanogaster* | (Chen et al., 2013) | Suppl. Fig. 8 |
| *Dmel Orco-Gal4* |  | BDSC_26818 | *D. melanogaster* | (Wang et al., 2003) | Figure 5,  Suppl. Fig. 8,  Suppl. Fig. 10 |
| *Dmel UAS-GCaMP6f* |  | BDSC_42747 | *D. melanogaster* | (Chen et al., 2013) | Figure 5,  Figure 6,  Suppl. Fig. 8,  Suppl. Fig. 9,  Suppl. Fig. 10,  Suppl. Fig. 11,  Suppl. Fig. 12 |
| *DsecOrco^Gal4^* |  |  | *D. sechellia* | (Auer et al., 2020) | Figure 5,  Suppl. Fig. 5,  Suppl. Fig. 8,  Suppl. Fig. 10 |
| *DsecOr22a^Gal4^* |  |  | *D. sechellia* | (Auer et al., 2020) | Figure 3,  Suppl. Fig. 5,  Suppl. Fig. 6, |
| *Dsec nSyb-ΦC31* | *nSyb-ΦC31 (Addgene #*133868) | *Dsec-attP26* (T.O.A. *unpublished*) | *D. sechellia* | attB/P integration, this study | Figure 3,  Suppl. Fig. 6 |
| *Dsec UAS-SPARC2-D-CsChrimson-Venus* | *pHD-Dsec-attP40-SPARC2-D-CsChrimson-Venus* | *D. sechellia* nanos-Cas9 | *D. sechellia* | CRISPR knock-in, this study | Figure 3 |
| *Dsec UAS-SPARC2-D-TNT-HA* | *pHD-Dsec-attP40-SPARC2-D-TNT-HA* | *D. sechellia* nanos-Cas9 | *D. sechellia* | CRISPR knock-in, this study | Figure 3,  Suppl. Fig. 6 |
| *Dmel Or22a/b^Gal4^* |  |  | *D. melanogaster* | (Chahda et al., 2019) | Suppl. Fig. 5 |
| *Dmel UAS-CsChrimson-Venus* |  | BDSC_55136 | *D. melanogaster* |  | Figure 3,  Suppl. Fig. 5 |
| *Dmel Or22a-Gal4* |  | BDSC_9952 | *D. melanogaster* |  | Figure 3,  Suppl. Fig. 6 |
| *Dmel nSyb-ΦC31* |  | BDSC_84151 | *D. melanogaster* |  | Suppl. Fig. 6 |
| *Dmel UAS-SPARC2-S-GFP* |  | BDSC_84148 | *D. melanogaster* |  | Suppl. Fig. 6 |
| *Dmel UAS-SPARC2-I-GFP* |  | BDSC_84147 | *D. melanogaster* |  | Suppl. Fig. 6 |
| *Dmel UAS-SPARC2-D-GFP* |  | BDSC_84146 | *D. melanogaster* |  | Suppl. Fig. 6 |

**Supplemental Table 3.** **Oligonucleotides used to generate single sgRNA expression vectors, *in situ* probe templates and *pBac* transgenesis vectors.**

Overhangs used for Gibson Assembly are shown in lowercase.

| **Target** | **Forward primer (5’-3’)** | **Reverse primer (5’-3’)** |
| --- | --- | --- |
| *DsimOr85b sgRNA* | GTCGCAAATAATCCAACGAAGCCC | AAACGGGCTTCGTTGGATTATTTG |
| *DsecOr47a  in situ probe* | ATGAAGCCAACGGAAATCCAAAAACCC | TTTCTGCACAAACCATTAAGAAGCTGC |
| *DsecOr88a  in situ probe* | ATGGAAAGTTTCCTCCAAGTACAG | GATGAGAAGGCCTCCATGTTTG |
| *pUAS-ChR2 CsChrimson* | atgccataggccacctattcgtcttcctacTGGGCGCGCCTAGTATGTAT | ccgagtctctgcactgaacattgtcagatcGGCCAGATCGATCCAGACAT |
| *pUC57(3xP3-DsRed)* | cataggccacctattcgtcttcctactgggATTACGCCAAGCTTGCATGC | tcagcgccctgcaccattatgttccggaggCCAGTGAATTCGAGCTCGGT |

**Supplemental Table 4.** **Oligonucleotides used to generate multi-sgRNA expression vectors.**

| **Target** | **Name** | **Sequence (5’-3’)** | **Resulting sgRNAs** |
| --- | --- | --- | --- |
| *Dseclz  & Dsimlz* | *PCR1fwd* | GCGGCCCGGGTTCGATTCCCGGCCGATGCAGAGGCGTCTGCTCTGCCGCCGTTTTAGAGCTAGAAATAGCAAG | sgRNA1: GAGGCGTCTGCTCTGCCGCC |
|  | *PCR1rev* | ATCGCTCCATCCACAATAGATGCACCAGCCGGGAATCGAACCC | sgRNA2: TCTATTGTGGATGGAGCGAT |
|  | *PCR2fwd* | TCTATTGTGGATGGAGCGATGTTTTAGAGCTAGAAATAGCAAG | sgRNA3: TAATAACAACGCCGTTCACC |
|  | *PCR2rev* | ATTTTAACTTGCTATTTCTAGCTCTAAAACGGTGAACGGCGTTGTTATTATGCACCAGCCGGGAATCGAACCC |  |
